# SWEET13 transport of sucrose, but not gibberellin, restores male fertility in Arabidopsis *sweet13;14*

**DOI:** 10.1101/2022.05.05.490848

**Authors:** Reika Isoda, Zoltan Palmai, Akira Yoshinari, Li Qing Chen, Florence Tama, Wolf B. Frommer, Masayoshi Nakamura

**Affiliations:** Institute of Transformative Bio-Molecules (WPI-ITbM), Nagoya University, Chikusa, Nagoya, 464-8601, Japan; Department of Plant Biology, University of Illinois at Urbana-Champaign, Urbana, Illinois, USA; Department of Physics, Graduate School of Science, Nagoya University, Aichi 464-8602, Japan; Center for Computational Science, RIKEN, Kobe, Hyogo 650-0047, Japan; Molecular Physiology, Heinrich-Heine-University, Düsseldorf, Germany

**Author notes:** **Authors for correspondence:** Masayoshi Nakamura and Wolf B. Frommer, **Email:**. **Author Contributions:** W.F. and M.N. designed the research; R.I. performed research; Z.P. performed molecular dynamics/molecular docking; L.Q. contributed plant materials; R.I., A.Y., F.T., M.N., and W.F. analyzed data; and R.I., M.N., and W.F. wrote the paper. R.I. and Z.P contributed equally to this work. **Competing Interest Statement:** The authors declare no competing interest.

**Keywords:** sucrose, gibberellin, transporter, promiscuity, male sterility

## Abstract

SWEET sucrose transporters play important roles in the allocation of sucrose in plants. Some SWEETs were shown to also mediate transport of the plant growth regulator gibberellin (GA). The close physiological relationship between sucrose and GA raised the questions of if there is a functional connection, and whether one or both of the substrates are physiologically relevant. To dissect these two activities, molecular dynamics were used to map the binding sites of sucrose and GA in the pore of SWEET13 and predicted binding interactions that might be selective for sucrose or GA. Transport assays confirmed these predictions. In transport assays, the N76Q mutant had 7x higher relative GA_3_ activity, and the S142N mutant only transported sucrose. The impaired pollen viability and germination in *sweet13;14* double mutants were complemented by the sucrose-selective SWEET13^S142N^ but not by the SWEET13^N76Q^ mutant, indicating that sucrose is the physiologically relevant substrate and that GA transport capacity is dispensable in the context of male fertility. Therefore, GA supplementation to counter male sterility may act indirectly via stimulating sucrose supply in male sterile mutants. These findings are also relevant in the context of the role of SWEETs in pathogen susceptibility.

## Introduction

We typically assign binary substrate-enzyme or substrate-transporter relationships. Transporters and enzymes are named based on a single assay or on selectivity towards a small number of related compounds. This concept is mainly based on limited test capacity. It is however now widely accepted that enzymes (and likely transporters) are promiscuous and can act on a much wider spectrum of substrates (1), likely many more than we currently conceive. Promiscuity is likely of importance for the efficacy of drugs and is relevant in the context of evolution. One may hypothesize that several substrates may be physiologically relevant, while many others may not currently be important for the function of the organism, making them irrelevant side activities. The relationship of kinetic properties and physiological concentrations of the substrates determine which substrate is effectively converted or transported.

Transporters are proteins that mediate the movement of solutes across the cell membrane, facilitating nutrition, signal transduction, cell-to-cell communication, and drug uptake and efflux. Highly sensitive assays have been able to identify plant amino acid transporters including NTR1, which in heterologous assays had a weak activity (2). Shortly afterwards, the first oligopeptide transporters were identified in yeast and mammals, which were strikingly similar to NTR1 (3-5). Functional assays validated that NTR1 functions as an oligopeptide transporter, indicating that amino acids most likely represent a non-relevant side activity (6). It was surprising that this peptide transporter was closely related to the plant nitrate transporters, with highly diverse substrate sizes. Notably, human PepT oligopeptide transporters are highly promiscuous and transport a wide spectrum of xenobiotic compounds, i.e., drugs (7). In plants, it was found that individual transporters in the nitrate/peptide transporter family have distinct, complex, overlapping activities for nitrate, chloride, oligopeptides, and glucosinolates, as well as most phytohormones including auxin, indole-3-butyric acid, abscisic acid, gibberellin (GA), and jasmonic acid conjugates (8, 9).

SWEETs represent a relatively recently described class of small transporters that mediate uniport of sugars (10, 11). The selectivity of SWEETs for sucrose and hexoses is correlated with phylogeny, for instance Clade 3 SWEETs mediate transport of sucrose. Physiologically, SWEETs fulfill key functions in nectar secretion, phloem loading, pollen nutrition, and seed filling that, at first sight, all appear to be consistent with sugar transport by SWEETs. However, Clade 3 SWEETs from *Arabidopsis* were shown to also mediate GA_3_ and GA_4_ uptake in a GA-dependent yeast three-hybrid (Y3H) assay (12). Surprisingly, GA transport capability was not conserved in Clade 3 SWEETs of rice (13). The use of stronger promoters allowed us to detect possibly a very weak GA-transport activity for OsSWEET11a in Y3H assays (14). *SWEET13* (*RPG2, Ruptured Pollen Grain2*), is, together with *SWEET8* (*RPG1*), essential for male fertility in *Arabidopsis* (15). SWEET13 and SWEET14 double mutants were also male sterile (12) and have been hypothesized to transport GA during anther dehiscence and subsequent seed formation. The restoration of male sterility by external application of GA was interpreted as strong evidence for a physiological role of the GA transport activity of SWEETs sterile (12). However, GA and sugars are tightly interconnected as GA affects growth and is apparently a key form for transport of reduced carbon and energy and GA and sugars have synergistic effects in affecting anther dehiscence and the release of viable pollen grains (16, 17). Therefore, the relative role of GA and sucrose transport activities could not be disentangled unambiguously. Notably, male sterility observed for the rice *ossweet11a;b* double mutant could not be rescued by GA supplementation (14). Therefore, it remains unknown if the dual selectivity of SWEETs is physiologically relevant and how it can mechanistically explain the parallel transport of such diverse compounds that also have different roles. In particular, phytohormone transport needs to be controlled tightly.

While GA_3_ and GA_4_ are very similar, differing only in the presence or absence of a hydroxyl group at position 13, the sucrose molecule, to our naïve eyes, appears structurally very different. The finding of complex, multi-substrate recognition in transporters raises many questions – are there multiple binding pockets and/or transport routes? How exactly are two apparently very diverse substrates recognized? What is the transport mechanism and directionality if one substrate (sucrose) is uncharged, while the other substrate carries a charge (GA anion)? Are mutant phenotypes due to defects in sucrose or GA transport deficiencies or both?

A key question was therefore whether one or both of the transport activities of the SWEETs are physiologically relevant. The most effective way to evaluate the relative roles of sucrose and GA transport activities would be to alter the relative selectivity of the SWEET, and to test whether more selective SWEETs are able to restore phenotypes of *sweet* knockout mutants. The availability of high-resolution structures of SWEET13 [Protein Data Bank (PDB) ID: 5XPD] (18) and site-directed mutagenesis in combination with transport assays enabled validation of the predictions. Notably, a more sucrose-selective transporter was able to restore pollen viability and germination defects of a *sweet13;14* double *knockout* mutants, while the more GA-selective variant was unable to correct either of the defects.

## Results

### Prediction of sucrose and GA interactions with SWEET13

The capability of SWEET13 to transport both sucrose and gibberellin had been demonstrated by different assays using yeast and animal cells (11, 12). As a first step for determining the relative role of the two transport activities, ligand binding was predicted using docking and MD simulations based on the structure of SWEET13 (PDB ID: 5XPD) (18). Since the 5XPD structure was obtained with a SWEET13 variant that was made more thermostable by mutagenesis, molecular docking was performed using a homology model in which the stabilizing mutations in SWEET13 were reverted to the native sequence and by removing the fused rubredoxin (see *SI Appendix*, Fig. S1).

The conformation of the binding sites for sucrose was grouped into three main clusters, while for GA_3_ only a single stable cluster was observed, which notably overlapped with Cluster 3 for sucrose (Fig. 1 A and B). The conformational subspace exploited by sucrose was thus substantially larger compared to GA_3_. In the most populated cluster, Cluster 1, sucrose occupied a vertical position along the channel axis (Fig. 1A; Table 1). The fructosyl moiety pointed toward the extracellular gate, making hydrogen bonds with Asn76 and Asn196 and hydrophobic contacts with Trp58 and Trp180. The glucosyl moiety pointed towards the intracellular gate, having hydrophobic interactions with Trp58, Trp180, Val23, and Val145. In Cluster 2, sucrose posed horizontally relative to the channel axis. The fructosyl unit made hydrophobic contacts with Trp58 and Trp180. The hydrogen of the glucosyl moiety bonded to Ser54 and Asn196 and made hydrophobic interactions with Trp58, Trp180, and Val145 (Fig. 1A; Table 1). The representative pose of sucrose in the least populated Cluster 3 was vertical, as in Cluster 1, with the glucosyl and fructosyl subunits pointing in- and outward, respectively (Fig. 1A; Table 1). The major difference between Clusters 1 and 3 was a 180-degree rotation of the glucosyl moiety around the glycosidic linkage. The fructosyl unit hydrogen bonded with Asn196 and made hydrophobic contacts with Trp58, Trp180, while the glucosyl unit hydrogen bonded with Asn76 and had hydrophobic contacts with Trp58, Trp180, and Val23.

**Figure 1.**
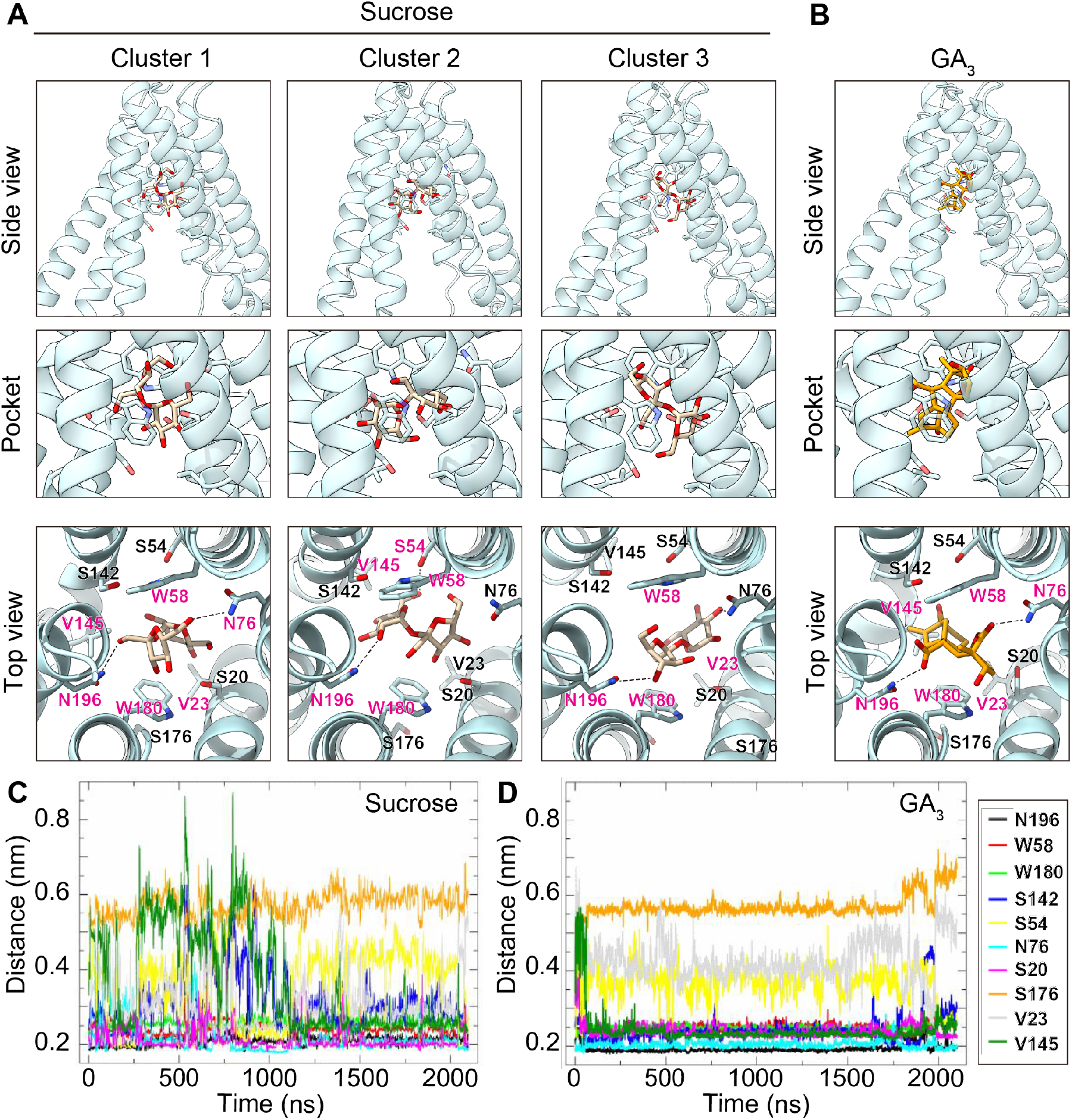
Molecular docking of sucrose and GA_3_ in SWEET13. Clusters of binding pocket conformations for SWEET13*Suc and SWEET13*GA_3_. Representative conformations of sucrose binding pose for (*A*) Clusters 1, 2, and 3, and (*B*) the binding pose for GA_3_. The color pink was used to indicate amino acids that provide hydrogen bonds and hydrophobic contacts with sucrose and GA_3_ (Table 1) (*C*) Minimum distance between sucrose and surrounding residues as a function of time. (*D*) Minimum distance between GA_3_ and surrounding residues as a function of time.

**Table 1.** Predicted interactions of SWEET13 binding pore amino acids with sucrose and GA_3_

| Side chain | Contact sucrose Cluster1 | Contact sucrose Cluster2 | Contact sucrose Cluster3 | Sucrose transport* | Contact GA<sub>3</sub> | Mutagenesis in this study |
| --- | --- | --- | --- | --- | --- | --- |
| S20 |  |  |  |  |  | S20A, S20N, S20Q |
| V23 | x |  | x | V23L: ~50% | x | V23A |
| S54 |  | x |  |  |  | S54A |
| W58 | x | x | x | W58A: ~50% | x |  |
| N76 | x |  |  | S76A: ~10% | x | N76Q |
| S142 |  |  |  | S142A: ~85% |  | S142A, S142L, S142N |
| V145 | x | x |  | V145A: ~50%<br>V145M: ~10% | x | V145A, V145L |
| W180 | x | x | x | W180A: ~10% | x |  |
| N196 | x | x | x |  | x | N196A, N196Q |

521 \* data from Han et al., 2017

The sole primary binding position of GA_3_ was vertical and established hydrogen bonds to Asn76 and Asn196 and hydrophobic contacts with Val23, Trp58, Val145, and Trp180 (Fig. 1B; Table 1). The conformational flexibility of sucrose relative to GA_3_ may be explained merely by ligand topology. In the case of sucrose, the possibility of torsional rotations around the glycosidic linkage likely allows for a greater number of binding modes. GA_3_ lacks a torsional linkage and is a larger molecule, which might be explain the tightly packed GA_3_ binding mode.

To gain a more precise knowledge of the ligand interactions, the minimal distances between the ligands and proposed binding residues were analyzed over time throughout the simulations. Sucrose maintained stable hydrogen bonds with Asn196, Asn76, and Ser20 and its hydrophobic contacts with Trp58 and Trp180 (Fig. 1C). In addition, labile interactions between sucrose and Ser142, Ser54, Val23, and Val145 were detected. Sucrose did not appear to interact with Ser176. In comparison, GA_3_ made tight and stable contacts with all residues involved in sucrose binding, except for Val23 and Ser54. GA_3_ was well anchored and rigidly held in place by its putative residues, establishing one primary binding mode, in contrast with the floppier positioning of sucrose in the binding pocket (Fig. 1D). Based on the simulation, we hypothesize that GA_3_ forms more stable interactions with Ser142 and Val145 relative to sucrose. These observations are consistent with the relatively low affinity of SWEETs for sucrose (in the millimolar range) and indicate that sucrose and GA transport will be competitive. Together, the docking models predicted that, while multiple side-chain interactions are shared by sucrose and GA_3_, Ser142/Val145 and Val23/Ser54 show preference for GA_3_ and sucrose, respectively, and may be targeted for mutagenesis to shift the relative selectivity for the two substrates (Table 1). Alanine substitution of Trp58, Asn76, Val145, Ser176, and Trp180 had previously been shown to be important for sucrose transport, and four residues, Val23, Ser54, Val145, and Ser176, were shown to be responsible for substrate selectivity between sucrose and glucose (11, 18). Of note, the substrate analog dCMP, which had been observed in the crystal structure of the thermostabilized SWEET13, takes the same vertical binding mode in Cluster 1 as sucrose and GA_3_ (*SI Appendix*, Fig. S2).

### Change in relative selectivity of SWEET13 by mutagenesis

To validate the predicted substrate interactions and the effect of individual side chains of SWEET13 on the relative selectivity towards sucrose and GA transport activities, thirteen mutations were introduced into SWEET13 by site-directed mutagenesis (Table 1). Sucrose and GA transport activities were quantified for each mutant using transport assays. Sucrose transport activity was quantified by co-expressing SWEET13 with the sucrose sensor FLIPsuc-90µΔ1 in mammalian HEK293T cells (11). Substitutions at N76, V145, and N196 to bulky residues (N76Q, V145L, and N196Q) caused substantially reduced sucrose transport activities (Fig. 2 A and B). Likewise, when S20 and S142 were replaced with bulky residues (S20N and S142L), sucrose transport activity was significantly reduced (Fig. 2 A and B). Although similarly bulky, S142N did not show a significant inhibition of sucrose transport. Replacing S20, V23, S54, S142, V145, and N196 with alanine did not substantially inhibit sucrose transport activity (*SI Appendix*, Fig. S3). The results indicate that bulky side chains in the binding pocket block the accessibility for sucrose, implicating both size and shape of the binding pocket in sucrose transport.

**Figure 2.**
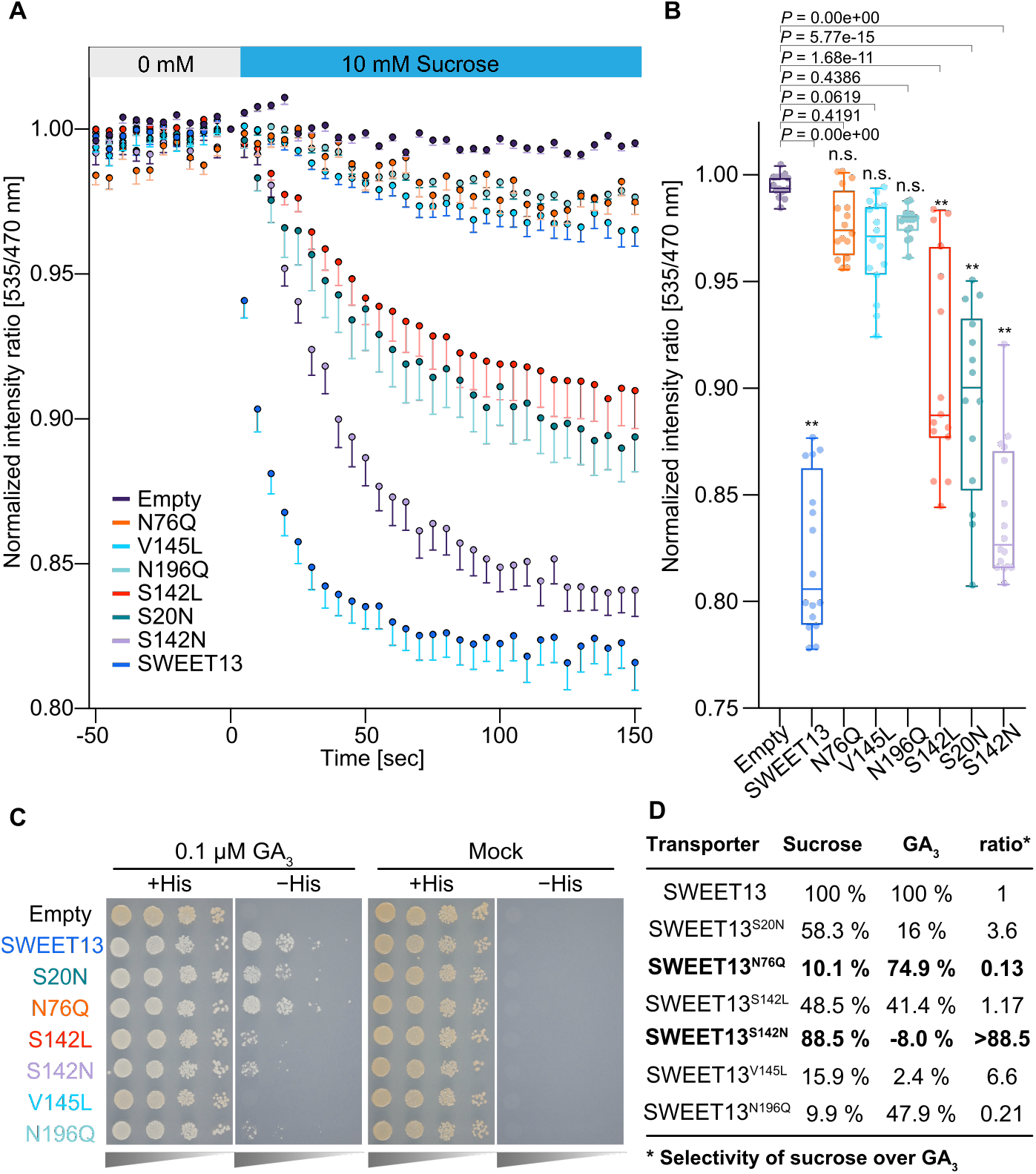
Sucrose and GA_3_ uptake assays of SWEET13 with mutations in proposed substrate binding sites. (A) Sucrose uptake activity in HEK293T cells expressing sucrose biosensor FLIPsuc-90µΔ1. Adding 10 mM sucrose led to negative ratio changes, consistent with sucrose accumulation in cells. In the absence of SWEET13 (Empty vector), no significant change in ratio was observed. Mutations in the binding sites of SWEET13 were tested (mean - s.e.m., n ≥ 14 cells). (B) Boxplot of average emission ratios from 125 to 150 s after addition of sucrose from the data in (A). The asterisk indicates a significant difference compared to empty vector (\*\**P* < 0.0001 by one-way ANOVA with Dunnett’s post-hoc test; n.s. indicates no significant difference). In the box plots, the box represents the range from the 25^th^ to 75^th^ percentile, each horizontal line marks the median value and the whiskers from the 2.5 to 97.5^th^ percentile. (C) Ten-fold serial dilution assay using yeast carrying both pDEST22-GID1a and pDEST32-GAI and either pDRf1-SWEET13, pDRf1-SWEET13 with mutations (S20N, N76Q, S142L, S142N, S145L, N196Q), or empty vector (negative control). Yeast cells were grown on SD (-Leu, -Trp, -Ura) or selective SD (-Leu, -Trp, -Ura, -His) medium, containing 3 mM 3-AT and 0.001% (v/v) DMSO as mock or 0.1 µM GA_3_, for 3 days at 30 °C. Comparable results were obtained in three independent replicates. (D) Sucrose and GA_3_ transport efficiency of SWEET mutants. The transporter activities for sucrose and GA_3_ were calculated from the results of uptake assays using HEK293T cells in Fig. 2B and Fig. S4, respectively.

Two independent transport assay systems, namely a GA-dependent yeast three-hybrid (Y3H) assay and the GA biosensor GPS1 in HEK293T cells, were used to measure GA transport activity of co-expressed SWEET13 variants (13, 19). In the presence of GA_3_, yeast expressing SWEET13^S142N/L^ grew less compared with wild-type SWEET13, while SWEET13^N76Q^ or SWEET13^N196Q^ restored the growth of yeast on the selection media (-His+GA_3_) (Fig. 2C). V145L did not show substantial GA or sucrose transport activity (Fig. 2B). To independently validate the GA transport activity, SWEET13 variants were expressed in HEK293T cells expressing the sensors GPS1. The YFP/CFP fluorescence ratio of GPS1 before and 3 hours after application of 1 µM GA_3_ was significantly higher in the presence of SWEET13. Mutants S20N/Q, S142A/L/N, V145L, and N196A showed significantly lower GA transport activity relative to wild type SWEET13 (*SI Appendix*, Fig. S4). Together, these results indicate that SWEET13^N76Q^ and SWEET13^N196Q^ preferentially transport GA_3_ over sucrose, whereas SWEET13^S142N^ preferentially transports sucrose over GA_3_ (Fig. 2D). Since the substitution of N196Q reduced GA transport activity of SWEET13 relative to N76Q, we used SWEET13^N76Q^ to validate the effect of a SWEET transporter with increased GA selectivity.

### Male fertility depends on sucrose transport activity of SWEET13

The *sweet13; sweet14* mutants has reduced fertility and fewer seeds in siliques compared to wild type, consistent with the combined role of SWEET13 and SWEET14 in male fertility (12). Since

GA_3_ application can restore fertility, one could hypothesize that only SWEET13^N76Q^, which retained GA_3_ transport but had reduced sucrose transport activity, might rescue male fertility of *sweet13; sweet14* mutants, while SWEET13^S142N^, which has reduced GA transport activity, but retained sucrose transport activity, would not. Complementation of the *sweet13; sweet14* mutant phenotype by SWEET13^N76Q^ and SWEET13^S142N^ was tested; wild-type SWEET13 served as a positive control. Pollen viability was evaluated by double staining with fluorescent diacetate (FDA; stains viable pollen with intact cell membranes) and propidium iodide (PI; stains pollen with compromised cell membrane) (20). In wild-type *Arabidopsis*, 97% of pollen grains were FDA positive, but only 8% of *sweet13; sweet14* pollen grains were stained (Fig. 3 A and B). Surprisingly, complementation with SWEET13^S142N^ (without GA transport) fully restored pollen viability of *sweet13; sweet14* mutants to the same level as wild-type SWEET13 (80%). In contrast, SWEET13^N76Q^ (without sucrose transport) did not lead to a significant increase in pollen viability of the *sweet13; sweet14* mutant (*P* = 0.215, one-way ANOVA with Dunnett’s post-hoc test). As an independent validation of pollen viability, pollen germination was analyzed and found to be consistent with the results of pollen viability tests (Fig 3 C and D). These observations intimate that the sucrose transport activity of SWEET13, but not its GA transport activity, is required for pollen viability and thus male fertility.

**Figure 3.**
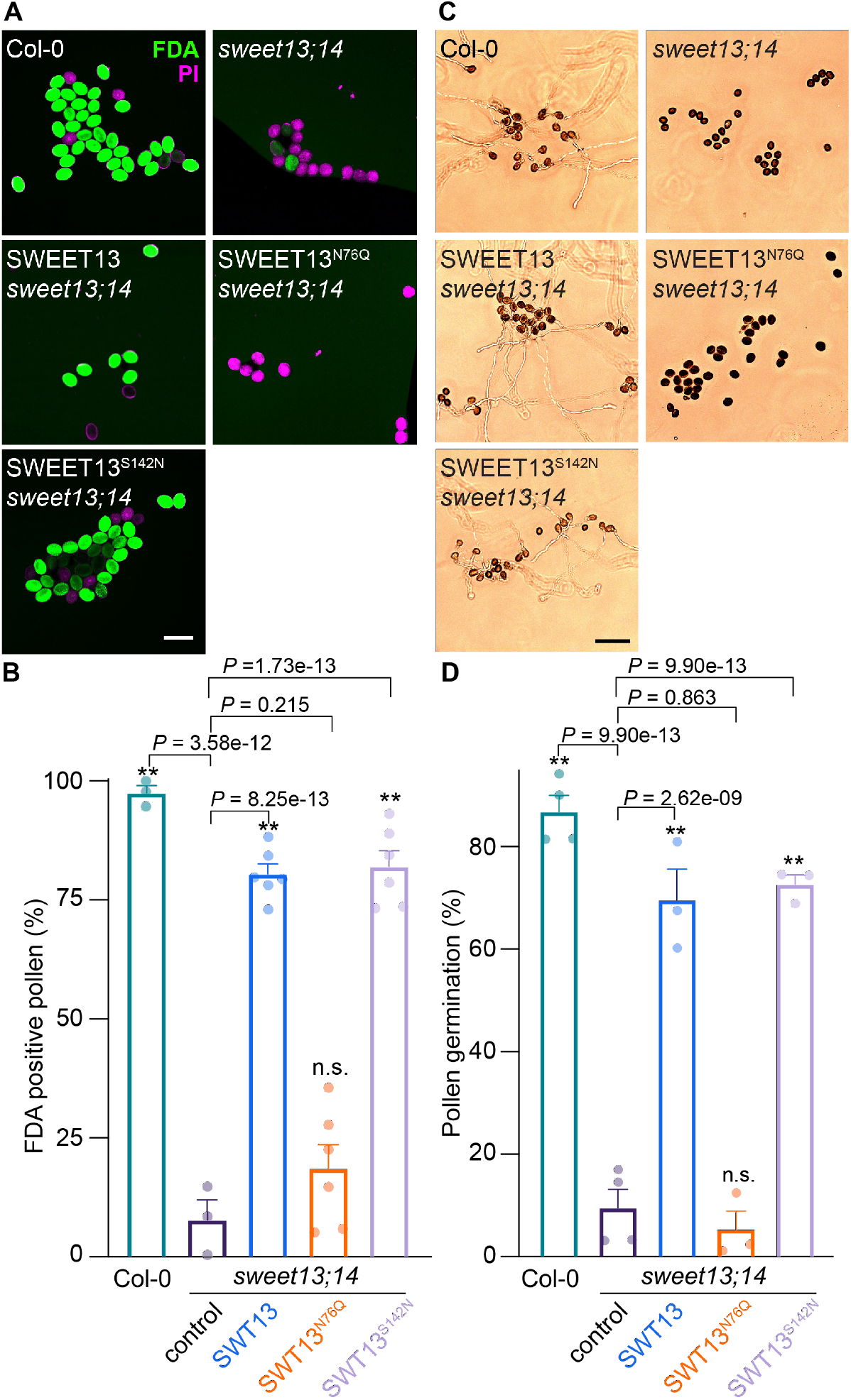
Restoration of pollen viability by the sucrose selective SWEET13^S142N^ variant. (A) Representative images of FDA/PI-stained pollen from wild type (Col-0), *sweet13; sweet14*, and *sweet13; sweet14* lines complemented with either SWEET13-GFP, SWEET13^N76Q^-GFP, or SWEET13^S142N^-GFP. Scale bar, 50 µm. (B) Quantification of the frequency of FDA-positive pollen in the respective lines. Data represent mean + SE from ≥3 independent assays with ≥100 pollen grains scored each time. (C) Representative images of pollen germination from the same lines. Scale bar: 100 µm. (D) Quantification of pollen germination frequency. Data represent mean + SE from ≥3 independent assay with ≥100 pollen grains each. *P-values shown* from one-way ANOVA with Dunnett’s post-hoc test. n.s., no significant difference.

### SWEET13 is produced in anther epidermis and endothecium

Recently, we discovered that Clade 3 SWEET members are involved in sugar transport-dependent male fertility in rice (14). OsSWEET11a and 11b accumulated in complementary sections of the vasculature of filament and anther, respectively (14). In contrast, *SWEET13 and 14* promoter-GUS reporter fusions did not appear active in the vasculature, but were in non-vascular tissues of the anther (12). To identify where the SWEET13 protein is expressed in *Arabidopsis* anthers, translational full-gene GFP fusions were investigated at different flower development stages. Of note, the SWEET13p-SWEET13-GFP fusion rescued the fertility of the *sweet13; sweet14* mutants (Fig. 3). GFP fluorescence was detectable only at late stages of anther development in the epidermis, endothecium and connective tissue in flowers at stages 13, thus emerging at a time just preceding dehiscence (Fig. 4 A-D; *SI Appendix*, Fig. S5). SWEET13 accumulation coincided with the third and highest wave of starch accumulation in anthers, in particular in the endothecium (21). SWEET13 was neither detected in the loculus nor in pollen grains (Fig. 4 B and C). No substantial accumulation of SWEET13p-SWEET13-GFP was found in anthers at the earlier stages, indicating that SWEET13 plays a specific role at late stages of anther development and pollen maturation. The data for protein accumulation shown here are consistent with the late accumulation of SWEET13 and 14 transcripts found in public microarray data (*SI Appendix*, Fig. S6). GFP fusions carrying the N76Q and S142N substitutions did not impact SWEET13 protein accumulation (*SI Appendix*, Fig. S7).

**Figure 4.**
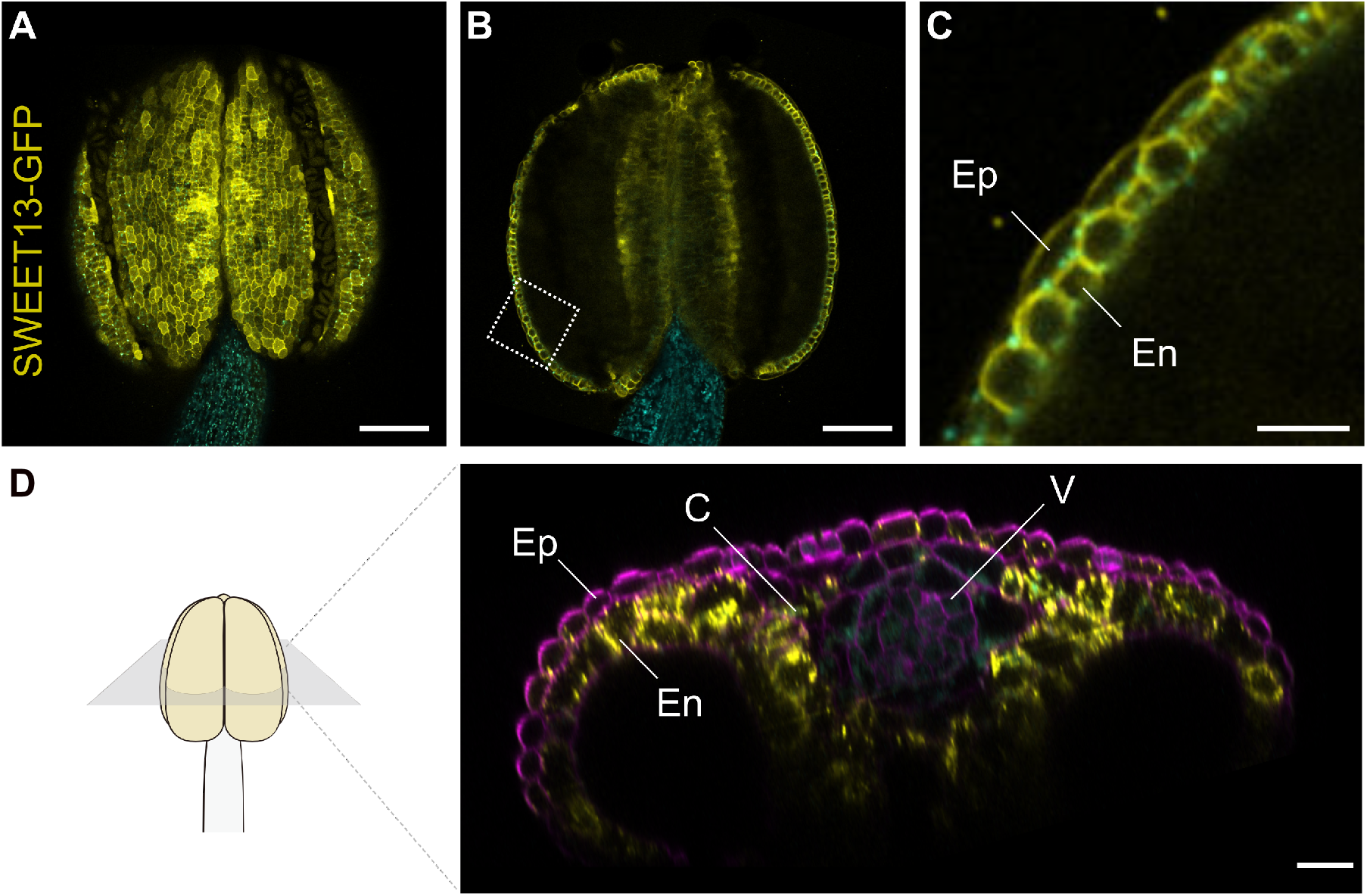
SWEET13 protein localization in Arabidopsis anthers. Confocal images of stage 13 anthers expressing *SWEET13-GFP* (yellow) in *sweet13; sweet14* at the (A) surface of anther, (B) middle section of anther; (C) enlarged view of boxed area in (B). (D) optical x/z section of stage 13 anther treated with ClearSee for 2 weeks and post-staining of cell wall with Calcofluor White (magenta). Cyan: autofluorescence. Abbreviations Ep, epidermis; En, endothecium; C, connective tissue; V, vasculature. Scale bars: 100 µm in (A) and (B), and 20 µm in (C) and (D).

## Discussion

Transporters, enzymes and receptors are not absolutely specific but can interact with multiple ligands, including xenobiotics. The interaction with xenobiotics is exploited for drug and pesticide development. The ability of transporters to recognize such compounds is relevant in the context of ADME/Tox, absorption, distribution, metabolism, elimination and toxicity (22). Recent studies had found that members of two plant transporter families – the NRT nitrate/PTR peptide transporters (also called NPF family, homologs of the mammalian PepT peptide transporter family; SLC15) and the SWEET sugar transporters – can also transport various plant hormones (8, 9, 12, 23). The promiscuity of PepTs is well established. Therefore, it is not be surprising that the plant homologs can transport as wide range of substrates. The physiological relevance of the ability to transport such a wide range of hormones remains to be elucidated. Here we tried to dissect the functional relevance of the sucrose and GA transport capabilities of the sucrose transporter SWEET13 *in vivo*.

Molecular docking performed here revealed that SWEET13 utilizes essentially the same cavity and binding pocket for GA_3_ and sucrose transport. Both substrates are predicted to hydrogen-bond with N76 and N196 (sucrose only in Cluster 1). When considering only the hydrogen bonds, both substrates could be similarly impacted by mutation. However, N76Q showed a more severe effect on sucrose transport activity, while GA transport was only slightly reduced. SWEET13^N76Q^ retained about 75% of its GA transport activity, while sucrose transport was reduced to 10%. As one may have expected from the smaller distance between GA and Ser142, SWEET13^S142N^ retained almost 90% of its sucrose transport activity, while GA transport activity was not detectable. The introduction of bulky side chains may result in steric hindrance of the tightly packed GA_3_, but not of the more flexible sucrose. These two mutants were thus used to evaluate the relative contribution of GA and sucrose to fertility. The full restoration of fertility by the sucrose transporting SWEET13^S142N^ but not by SWEET13^N76Q^ intimates that the GA transport function is dispensable for fertility.

By dissecting the selectivity of SWEET13 for sucrose and GA, we found that the sucrose transport activity, but not GA transport activity, is critical for pollen development and consequently male fertility. Physiological studies and localization analyses using GFP-fused SWEET13 indicated that SWEET13 functions in sucrose translocation from connective tissue to pollen grains. The sucrose transport by SWEET13 localized at the connective tissue and the epidermis and endothecium cells might therefore be important for pollen development.

GA application had been shown to compensate for the male fertility defect *in sweet13;14* double mutant plants (12). One potential explanation of the compensatory effect of external GA is the induction of metabolic processes that compensate for sucrose transport deficiency, either by altering SWEET gene expression or induction of cell wall invertases by GA, which has been observed (24). Alternatively, sucrose transport deficiency could be responsible for lower GA levels, and spraying GA circumvents this deficiency or the need for sucrose. We also tried to evaluate the restoration of the dehiscence defect of sweet13;14 double mutants, however the phenotype was highly variable, precluding reliable interpretation of the results.

The sucrose-transporting SWEETs show functional redundancy for male fertility in rice. An *ossweet11a; 11b* double *knockout* mutant was found to be male sterile (14). Mutant pollen was deficient in starch accumulation of pollen grains. Double *ossweet11a; 11b* mutants thus have similar phenotype as *atsweet13;14* mutants, which may be ascribed to orthologous functions, at first sight. Of note, the nomenclature of the SWEETs was based on phylogenetic analyses, however, Clade 3 SWEETs of *Arabidopsis* and rice do form separate subclades and thus were amplified after the ancestral split. OsSWEET11a and 11b are thus not orthologous to SWEET13 and 14 from *Arabidopsis* from a phylogenetic perspective. Moreover, OsSWEET11a and 11b proteins were detected in the veins of the anther, with perfectly complementary localization in the peduncle of the anther (OsSWEET11a), and the adjacent veins that enter the axis of anther (OsSWEET11b) (14). This pattern is distinct from that of SWEET13 and 14 in *Arabidopsis*, as described here for a GFP and GUS fusions (12). At present, we cannot exclude that additional Clade 3 SWEETs are fulfilling the orthologous functions. Another major difference to *Arabidopsis* is that the external application of GA_3_ did not restore male fertility in rice (14).

A number of SWEETs that transport GAs have been identified here and in previous studies (12, 13). The study here found that replacing Ser142 of SWEET13 with Asn interfered with GA transport activity. Amino acid sequence alignment of *Arabidopsis* and rice SWEETs revealed that SWEETs determined to be GA transporters, such as Clade 3 family members and OsSWEET3a, predominantly have Ser142, and OsSWEETs unable to transport GAs have Gly rather than Ser in the site (*SI Appendix*, Fig. S8). SWEET13^S142A^, also showed reduced GA transport activity, indicating that this amino acid is of importance for GA transport. Other untested SWEETs with Ser142 amino acids, such as SWEET1 and SWEET3 in *Arabidopsis*, may also have GA transport capacity. Val145 was also conserved in SWEET proteins that transport GAs, however, V145 is known to be an amino acid that confers selectivity for sucrose and hexose (Han et al., 2017). Replacing this Val145 amino acid with Leu reduced sucrose transport activity as well as GA transport activity (Fig. 2D), indicating that V145 is responsible for both sucrose and GA transport activities of SWEETs.

### Contribution of SWEET13 in sucrose flow from anther to pollen

The link between pollen viability and sucrose supply to developing anthers via the phloem appears obvious. However, the process seems to be highly complex, involving SWEET and SUT/SUC transporters in the vasculature of the stamen as well as the different cell types of the anthers and the microspores/pollen grains during multiple phases. There are three consecutive starch accumulation/degradation phases during anther and pollen development (21). Starch first accumulates in filaments and connective tissue, but not in the microspores (floral stages 5-8). While microspore development proceeds to pollen mitosis I, a second wave of starch accumulation is initiated in anthers and pollen (floral stages 10-12). Starch deposits become visible in the bicellular pollen grains and anther simultaneously. The pollen grains undergoing pollen mitosis II reach maximal starch accumulation, while starch is rapidly degraded in the anther. Soon afterwards, a third wave of starch accumulation occurs in the endothecium and connective tissue (floral stages 12-13). *SUC1* and *SUC2* H^+^/sucrose symporters are produced in these stages (floral stage 12-13) in the connective tissue and the filament, respectively (17, 25). Accumulation of SWEET13 protein in both epidermis and endothecium coincides with the third wave of starch accumulation in the anthers (floral stage 13). Sucrose is delivered by the phloem to the anthers. GFP, expressed using the AtSUC2 promoter, is confined to the phloem of the filament and symplasmically unloads sucrose into the anther connective tissue, from where it must be transported toward the anther locules (25). Sequential cycles of sucrose hydrolysis, starch biosynthesis and starch mobilization then enable efficient and temporally controlled sugar supply to the locule prior to feeding of pollen grains. SUC1 and SUC2, as secondary active sucrose transporters in stages 12-13, and SWEET13/14, in stage 13, thus contribute to specific cellular import and export steps. Amylolysis during the third wave may cause an increase in endothecium sucrose concentrations, leading to sucrose efflux to the apoplasm by SWEET13 down a concentration gradient into the locule. Involvement of hexose transporting SWEETs, such as SWEET8 (RPG1), indicates that there are additional steps involving apoplasmic invertases and the transport of hexoses, as in the case of maize kernel filling (26, 27).

GA and sucrose functions are tightly interrelated and both are required for growth, therefore one may hypothesize that GA supplementation rescued the fertility of the *sweet13;14* double mutant, though an indirect effect. Of note, many physiological studies have implied that Clade 3 SWEETs function in sucrose transport processes, and in all cases starch levels were impacted, including phloem loading in Arabidopsis and maize, seed filling in Arabidopsis and rice, and, notably, nectar secretion in Arabidopsis (28-32). In the case of phloem loading, the presence of SWEET11, 12 and 13 in the phloem parenchyma – the cell predicted to supply sucrose to the adjacent sieve element companion cell complex, which imports sucrose via the SUT H^+^/sucrose symporter – provides additional support for the physiological function in sucrose transport (33). Since Clade 3 SWEETs are hijacked by the rice blight pathogen *Xanthomonas oryzae pv. oryzae* (*Xoo*), it has been proposed that SWEET-mediated release of sucrose supports effective reproduction and thus virulence of *Xoo* in the xylem (34, 35). The recently revealed role of the sucrose utilization gene cluster in virulence may provide another piece of evidence showing that sucrose is the relevant substrate (36).

This work may also provide a roadmap for dissecting how nitrate, oligopeptides and plant growth regulators are transported by members of the nitrate/peptide transporter family, e.g., auxin by NTR1;1, or ABA, GA and JA by AIT1 and 3 (8, 9, 23, 37), and what this transport influences. This work also may aid in development of pesticides and antimicrobials for crop protection.

## Materials and Methods

### Sucrose transport assay in mammalian cells

For sucrose transport assays, HEK293T cells were co-transfected with constructs carrying the sucrose sensor FLIPsuc-90µΔ1 (38) and SWEET13 variants. Experimental details are described in *SI Appendix, SI Text*.

### GA transport assays in a yeast three-hybrid system

GA transport assays using a Y3H system (13, 14) are described in *SI Appendix, SI Text*.

### Plant materials

The *sweet13;14* double mutant was obtained by crossing *sweet13* (SALK_087791) with *sweet14* (SALK_010224), the same mutants described by (12). Vector constructs for SWEET13-GFP, plant growth conditions, and confocal imaging are described in *SI Appendix, SI Text*.

### Pollen viability and *in vitro* pollen germination assays

Propidium iodide-Fluorescent Diacetates (PI-FDA) assays for pollen viability and *in vitro* pollen germination assays were performed as described (20, 39). Experimental details are provided in *SI Appendix, SI Text*.

## Data Availability

The docking models of SWEET13 with sucrose and GA have been deposited and will be available at https://www.modelarchive.org/.

## Acknowledgments

We thank Drs. Michael Wudick and Ji-Yun Kim for valuable advice. We are particularly grateful for the outstanding technical assistance by Yukie Kawase. This research was supported by grants from Deutsche Forschungsgemeinschaft (DFG, German Research Foundation) under Germany’
ss Excellence Strategy – EXC-2048/1 – project ID 390686111, the Alexander von Humboldt Professorship (WF), and a Japan Society for the Promotion of Science (JSPS) grant-19H00932 (WF). ITbM is supported by World Premier Institute Center Initiative (WPI), Japan.

## References

1. O. Khersonsky, D. S. Tawfik, Enzyme promiscuity: a mechanistic and evolutionary perspective. Annu. Rev. Biochem. 79, 471–505 (2010).

2. W. B. Frommer, S. Hummel, D. Rentsch, Cloning of an Arabidopsis histidine transporting protein related to nitrate and peptide transporters. FEBS letters 347, 185–189 (1994).

3. M. Boll, et al., Expression cloning of a cDNA from rabbit small intestine related to proton-coupled transport of peptides, ßlactam antibiotics and ACE-inhibitors. Pflugers Arch. 429, 146–149 (1994).

4. Y. J. Fei, et al., Expression cloning of a mammalian proton-coupled oligopeptide transporter. Nature 368, 563–566 (1994).

5. J. R. Perry, M. A. Basrai, H. Y. Steiner, F. Naider, J. M. Becker, Isolation and characterization of a Saccharomyces cerevisiae peptide transport gene. Mol. Cell. Biol. 14, 104–115 (1994).

6. D. Rentsch, et al., NTR1 encodes a high affinity oligopeptide transporter in Arabidopsis. FEBS letters 370, 264–268 (1995).

7. M. Killer, J. Wald, J. Pieprzyk, T. C. Marlovits, C. Löw, Structural snapshots of human PepT1 and PepT2 reveal mechanistic insights into substrate and drug transport across epithelial membranes. Sci. Adv 7, eabk3259 (2021).

8. M. E. Jørgensen, et al., Origin and evolution of transporter substrate specificity within the NPF family. eLife 6, 29931 (2017).

9. Y. Chiba, et al., Identification of Arabidopsis thaliana NRT1/PTR FAMILY (NPF) proteins capable of transporting plant hormones. J. Plant Res. 128, 679–686 (2015).

10. L.-Q. Chen, et al., Sugar transporters for intercellular exchange and nutrition of pathogens. Nature 468, 527–532 (2010).

11. L.-Q. Chen, et al., Sucrose efflux mediated by SWEET proteins as a key step for phloem transport. Science 335, 207–211 (2012).

12. Y. Kanno, et al., AtSWEET13 and AtSWEET14 regulate gibberellin-mediated physiological processes. Nat. Commun. 7, 13245 (2016).

13. M. Morii, et al., The dual function of OsSWEET3a as a gibberellin and glucose transporter ts important for young shoot development in rice. Plant Cell Physiol. 61, 1935–1945 (2020).

14. L.-B. Wu, et al., OsSWEET11b, a potential sixth leaf blight susceptibility gene involved in sugar transport-dependent male fertility. New Phytol. 234, 975–989 (2022).

15. M. -X. Sun, X.-Y. Huang, J. Yang, Y.-F. Guan, Z.-N. Yang, Arabidopsis RPG1 is important for primexine deposition and functions redundantly with RPG2 for plant fertility at the late reproductive stage. Plant Reprod. 26, 83–91 (2013).

16. T. Chhun, et al., Gibberellin regulates pollen viability and pollen tube growth in rice. Plant Cell 19, 3876–3888 (2007).

17. R. Stadler, E. Truernit, M. Gahrtz, N. Sauer, The AtSUC1 sucrose carrier may represent the osmotic driving force for anther dehiscence and pollen tube growth in Arabidopsis. Plant J. 19, 269–278 (1999).

18. L. Han, et al., Molecular mechanism of substrate recognition and transport by the AtSWEET13 sugar transporter. Proc. Natl. Acad. Sci. USA 114, 10089–10094 (2017).

19. A. Rizza, A. Walia, V. Lanquar, W. B. Frommer, A. M. Jones, In vivo gibberellin gradients visualized in rapidly elongating tissues. Nat. Plants 3, 803–813 (2017).

20. X. Zhou, et al., SYP72 interacts with the mechanosensitive channel MSL8 to protect pollen from hypoosmotic shock during hydration. Nat. Commun. 13, 73–14 (2022).

21. A. Hedhly, et al., Starch turnover and metabolism during flower and early embryo development. Plant Physiol. 172, 2388–2402 (2016).

22. J. Keogh, B. Hagenbuch, C. Rynn, B. Stieger, G. Nicholls, “Membrane Transporters: Fundamentals, Function and Their Role in ADME” in Drug Transporters: Volume 1: Role and Importance in ADME and Drug Development, (The Royal Society of Chemistry, 2016), pp. 1–56.

23. Y. Kanno, et al., Identification of an abscisic acid transporter by functional screening using the receptor complex as a sensor. Proc. Natl. Acad. Sci. USA 109, 9653–9658 (2012).

24. R. K. Proels, M.-C. González, T. Roitsch, Gibberellin-dependent induction of tomato extracellular invertase Lin7 is required for pollen development. Funct. Plant Biol 33, 547–554 (2006).

25. A. Imlau, E. Truernit, N. Sauer, Cell-to-cell and long-distance trafficking of the green fluorescent protein in the phloem and symplastic unloading of the protein into sink tissues. Plant Cell 11, 309–322 (1999).

26. L. Sun, et al., Sugar homeostasis mediated by cell wall invertase GRAIN INCOMPLETE FILLING 1 (GIF1) plays a role in pre-existing and induced defence in rice. Mol Plant Pathol 15, 161–173 (2014).

27. D. Sosso, et al., Seed filling in domesticated maize and rice depends on SWEET-mediated hexose transport. Nature genetics 47, 1489–1493 (2015).

28. C. Zhang, et al., Two evolutionarily duplicated domains individually and post-transcriptionally control SWEET expression for phloem transport. New Phytol. 232, 17993–1807 (2021).

29. I. W. Lin, et al., Nectar secretion requires sucrose phosphate synthases and the sugar transporter SWEET9. Nature 508, 546–549 (2014).

30. M. Bezrutczyk, et al., Impaired phloem loading in zmsweet13a,b,c sucrose transporter triple knock-out mutants in Zea mays. New Phytol. 218, 594–603 (2018).

31. J. Yang, D. Luo, B. Yang, W. B. Frommer, J.-S. Eom, SWEET11 and 15 as key players in seed filling in rice. New Phytol. 218, 604–615 (2018).

32. L. Ma, et al., Essential role of sugar transporter OsSWEET11 during the early stage of rice grain filling. Plant Cell Physiol. 58, 863–873 (2017).

33. J.-Y. Kim, et al., Distinct identities of leaf phloem cells revealed by single cell transcriptomics. Plant Cell 33, 511–530 (2021).

34. R. Oliva, et al., Broad-spectrum resistance to bacterial blight in rice using genome editing. Nat. Biotechnol. 37, 1344–1350 (2019).

35. J.-S. Eom, et al., Diagnostic kit for rice blight resistance. Nat. Biotechnol. 37, 1372–1379 (2019).

36. M. Sadoine, et al., Sucrose-dependence of sugar uptake, quorum sensing and virulence of the rice blight pathogen Xanthomonas oryzae pv. oryzae. bioRxiv, 2021.08.22.457195 (2022).

37. G. Krouk, et al., Nitrate-regulated auxin transport by NRT1.1 defines a mechanism for nutrient sensing in plants. Dev. Cell 18, 927–937 (2010).

38. I. Lager, L. L. Looger, M. Hilpert, S. Lalonde, W. B. Frommer, Conversion of a putative Agrobacterium sugar-binding protein into a FRET sensor with high selectivity for sucrose. J. Bio. Chem. 281, 30875–30883 (2006).

39. K. Muro, et al., ANTH domain-containing proteins are required for the pollen tube plasma membrane integrity via recycling ANXUR kinases. Commun Biol 1, 152 (2018).

