## Supplementary Data for "SWEET13 transport of sucrose, but not gibberellin, restores male fertility in Arabidopsis *sweet13;14*"

Dissection of the role of sucrose and gibberellin transport activities of SWEET13 in male fertility of Arabidopsis

**This PDF file includes:**

Figures S1 to S8

Tables S1

Supplementary information text

SI References

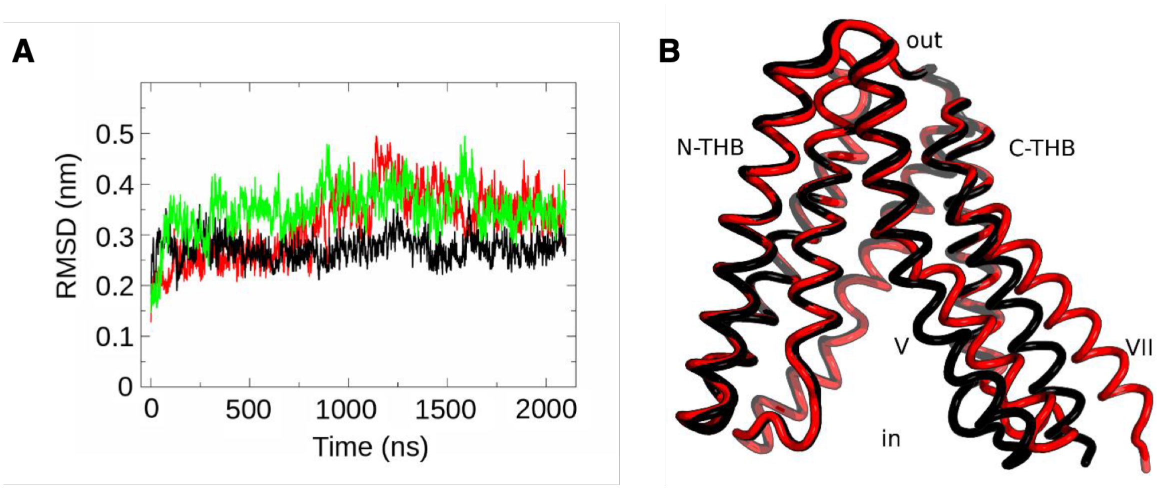


**Figure S1.** Time evolution of the RMSDs for apo and bound forms of SWEET13.

(A) Time evolution of the RMSDs compared to the initial coordinates (t=0 ns) for apo SWEET13 (black), and the models of SWEET13 bound to sucrose (SWEET13*Suc; red) and gibberellic acid (SWEET13*$GA_3$; green). (B) 3D structures of the SWEET13*Suc averaged for the first (0-900 ns, black cartoon) and second (900-2100 ns, red cartoon) part of the simulation.

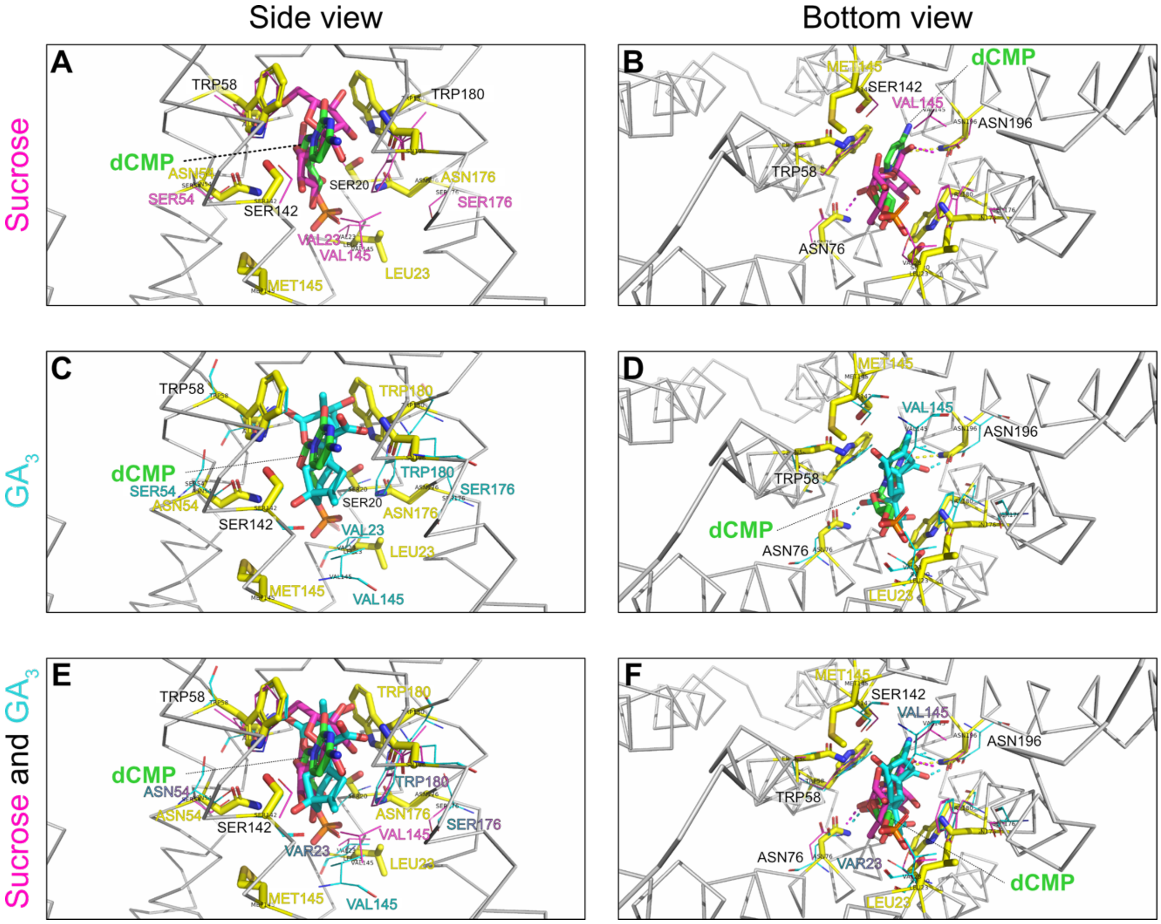


**Figure S2.** Comparison of docking clusters for dCMP, sucrose and $GA_3$

Representative conformation of (A) sucrose (magenta sticks) - cluster 1 binding mode - with its binding residues (magenta lines) in side and (B) bottom view, (C) $GA_3$ (cyan sticks) - cluster 1 binding mode - with its binding residues (cyan lines) in side view and D) bottom view, E) sucrose (magenta sticks) / $GA_3$ (cyan sticks) - cluster 1 binding mode - with their binding residues (magenta/cyan lines, correspondingly) in side view and in F) bottom view, superimposed onto the carbon alpha atoms of the 5XPD crystal structure (gray ribbon) with dCMP bound in the central cavity (green sticks). The backbone is shown only for the 5XPD crystal structure for the sake of clarity. The dCMP binding residues are shown as yellow sticks.

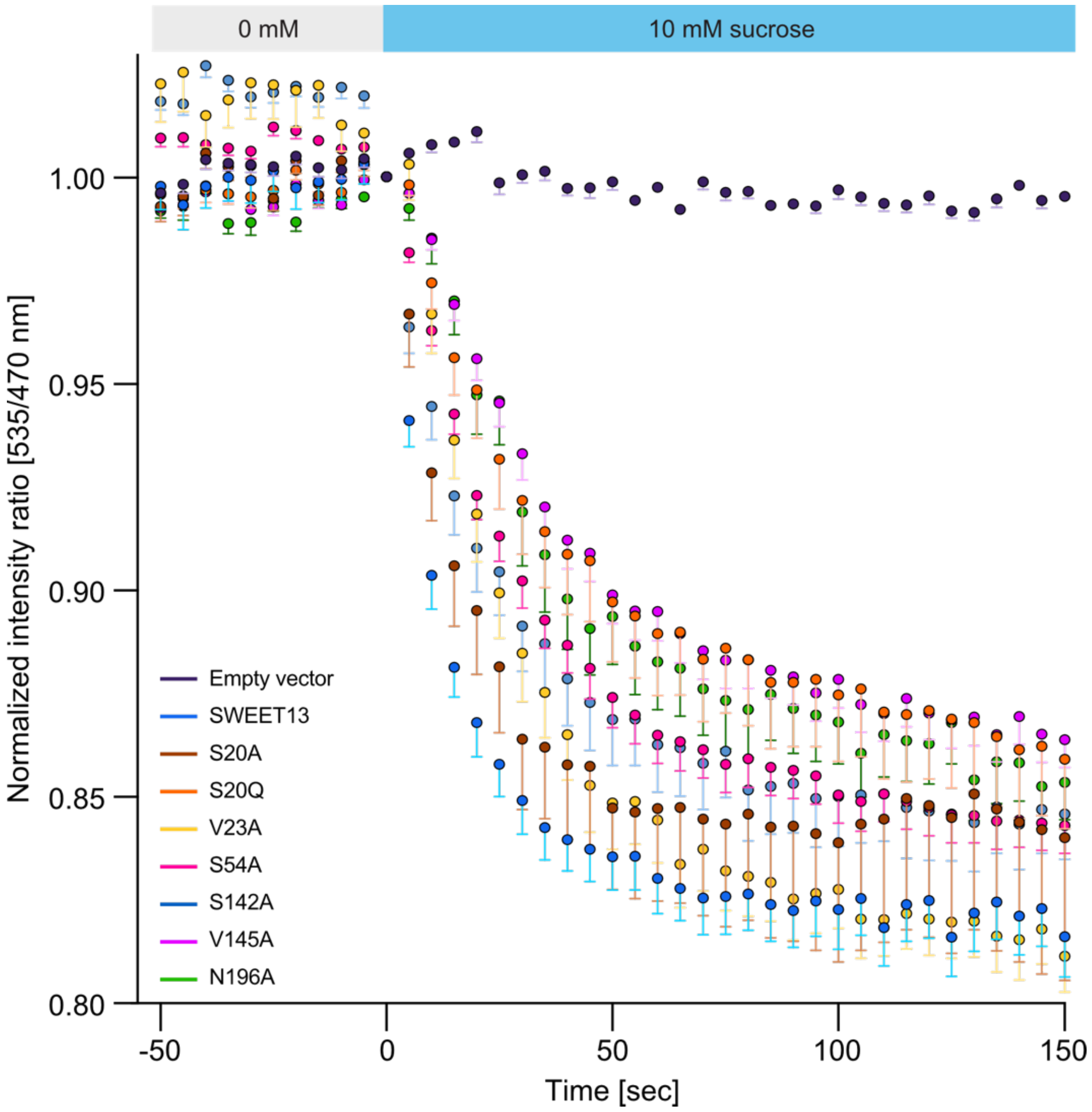


**Fig. S3.** Sucrose uptake by SWEET13 variants carrying mutations in the proposed substrate-binding pocket

Sucrose uptake activity was determined in HEK293T cells coexpressing the sucrose sensor FLIPsuc-90µΔ1 with SWEET13 or one of seven variants carrying mutations in the proposed binding site. Addition of 10 mM sucrose triggered a time-dependent negative ratio change, consistent with the accumulation of sucrose in the cells. In the absence of SWEET13 (Empty vector), no significant change in ratio was observed. (mean – s.e.m., n ≥ 14 cells). The experiment was repeated 3 times with comparable results.

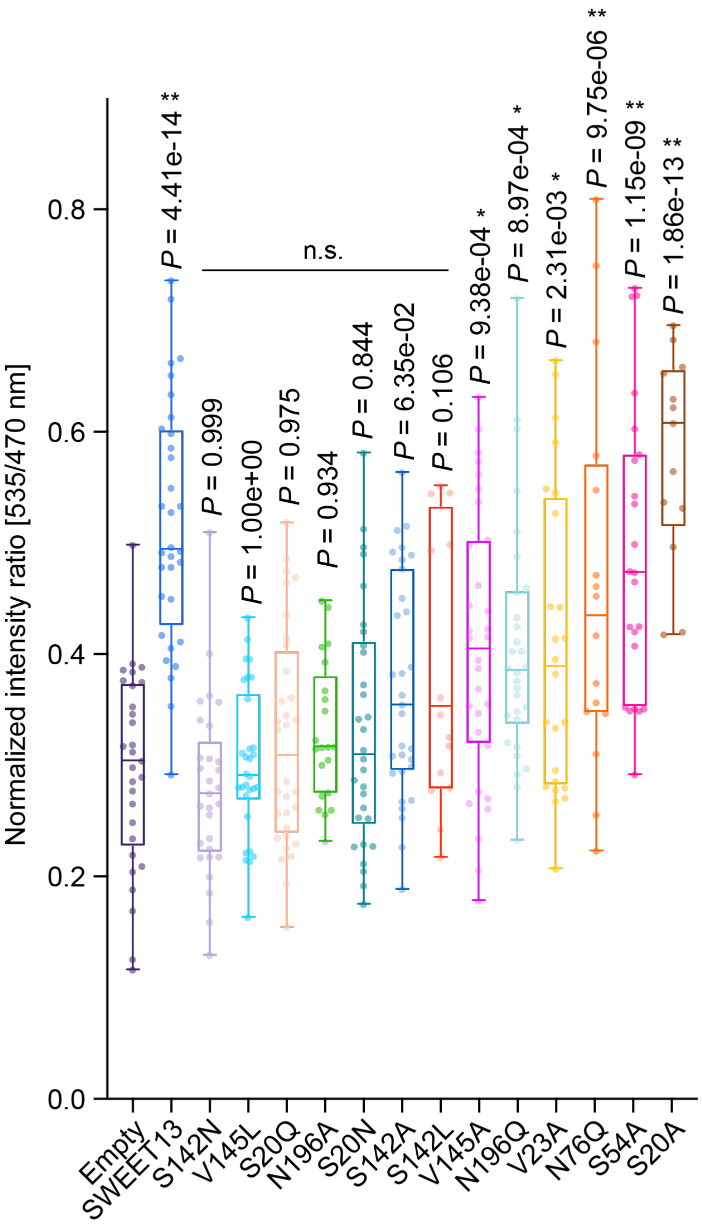


**Fig. S4.** $GA_3$ uptake by SWEET13 variants carrying mutations in the proposed substrate binding pocket

Boxplot of $GA_3$ uptake assays in HEK293T cells coexpressing the GA sensor GPS1 and SWEET13 (n =32 cells) or SWEET13 with mutations such as S142N (n = 29), V145L (n = 30), S20Q (n = 32), N196A (n = 21), S20N (n = 20), S142A (n = 31), S142L (n = 12), V145A (n = 32), N196Q (n = 30), V23A (n = 24), N76Q (n = 16), S54A (n = 23), S20N (n = 13). Empty vector served as negative control (n = 29). The YFP/CFP fluorescence ratio of GPS1 was measured after incubation with 1 µM $GA_3$ for 3 h. Asterisk indicates significant difference compared to 'Empty vector' control (**$P < 0.0001$, *$P < 0.01$ by one-way ANOVA with Dunnett's post hoc test). n.s.: no significant difference. In box plots, box represents the range from 25$^{th}$ to 75$^{th}$ percentile, the horizontal line marks the median value, and the whiskers from the 2.5 to 97.5 percentile. Each data point shown was generated by determining the mean emission ratio during 3-minute recordings. The experiment was repeated 3 times with comparable results.

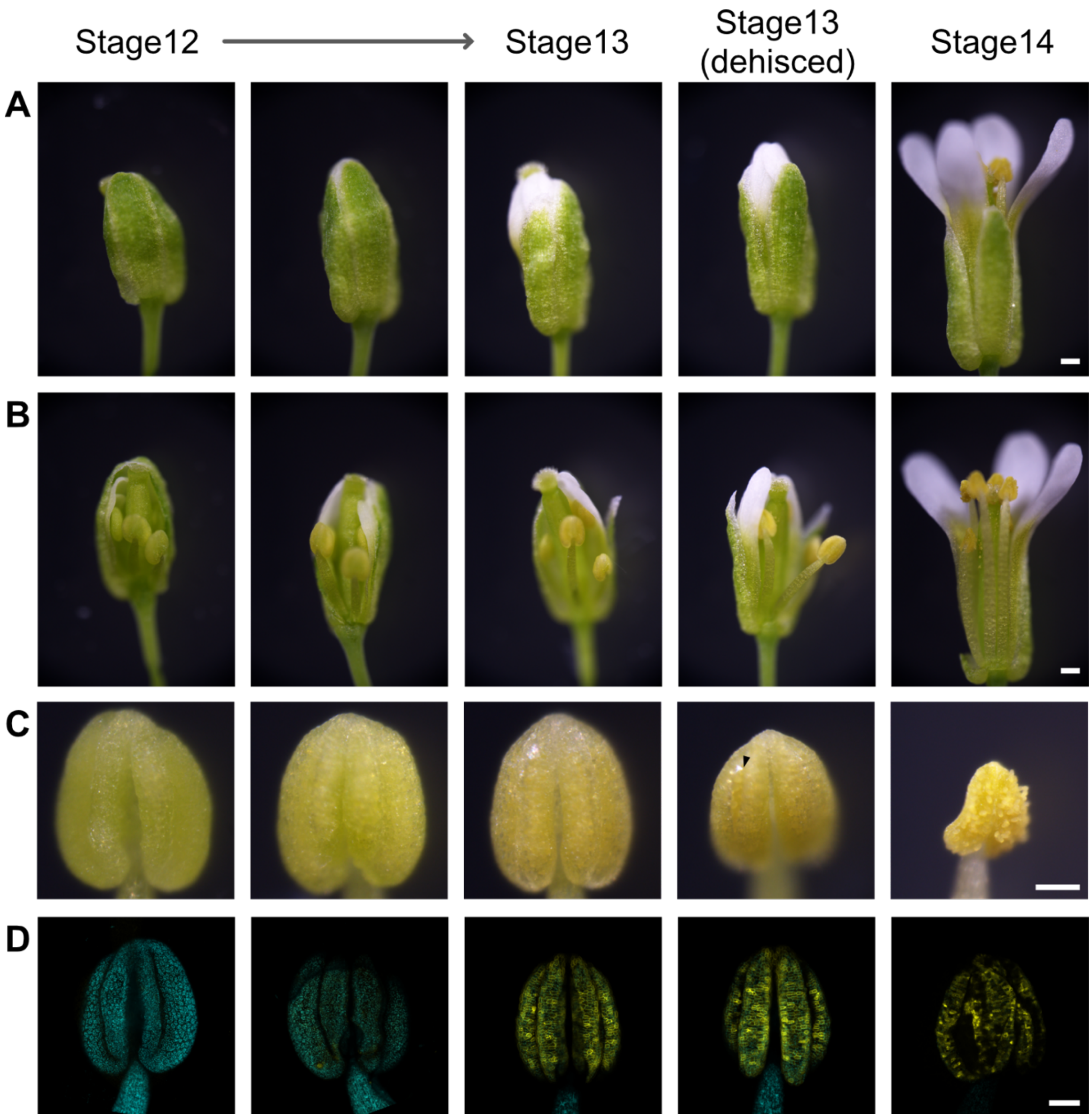


**Fig. S5.** SWEET13 protein accumulation in anthers at different stages of flower development

Phenotypes and GFP fluorescence at stages from 12 to 14 for *sweet13; sweet14* mutants expressing translational *SWEET13-GFP* fusions driven from the SWEET13 promoter in (A) buds/flowers, (B) pistil and stamen (C) anther, and (D) Confocal fluorescence images of sum slices projection of confocal images of anthers (yellow). Cyan indicates autofluorescence. Arrowhead indicates dehiscence. Scale bars: 200 µm in (A) and (B), and 100 µm in (C) and (D). Comparable results were obtained in 3 independent analyses.

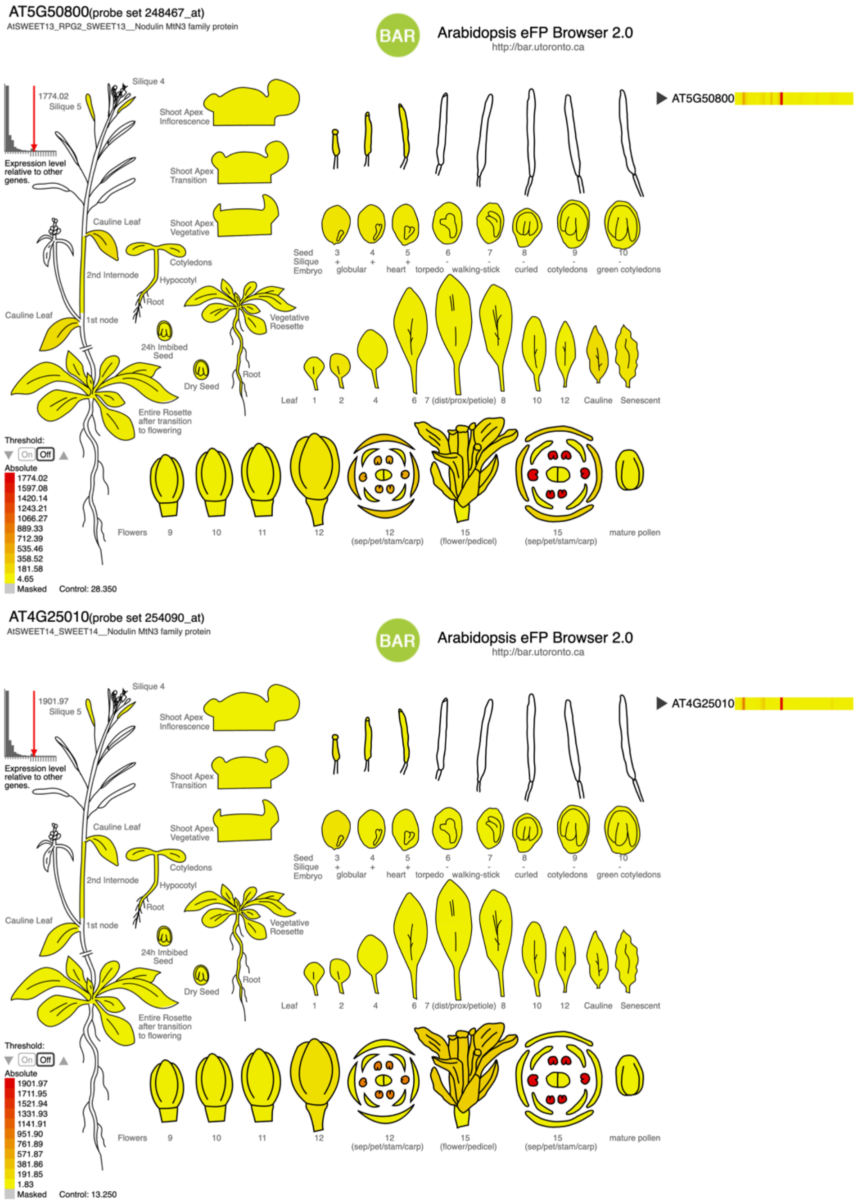


**Fig. S6.** Tissue specificity of SWEET13 and SWEET14 mRNA levels

SWEET13 (AT5G50800), SWEET14 (AT4g25010) expression data from database (Arabidopsis eFP Browser 2.0).

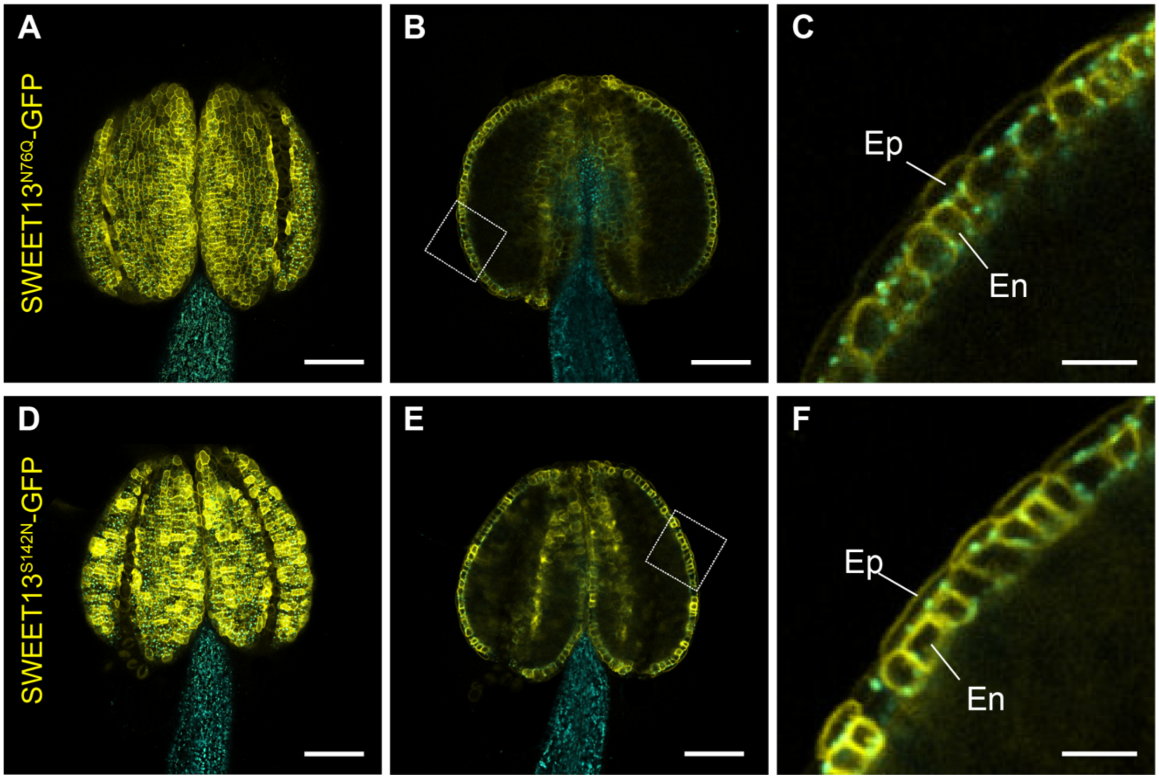


**Fig. S7.** Accumulation of the mutated versions of SWEET13 protein in *Arabidopsis* anthers

Confocal images of stage 13 anther expressing (A, B, C) translational *SWEET13*$^{N76Q}$*-GFP* and (D, E, F) *SWEET13*$^{S142N}$*-GFP* fusions driven from the SWEET13 promoter in *sweet13; sweet14* mutants. (A, D) surface of anther, (B, E) middle section of anther, and (C, F) enlarged view of boxed area in (B, E). Cyan indicates autofluorescence. Ep, epidermis; En, endothecium. Scale bars: 100 µm in (A, B, D, E), and 20 µm in (C, F). Comparable results were obtained in 3 independent analyses.

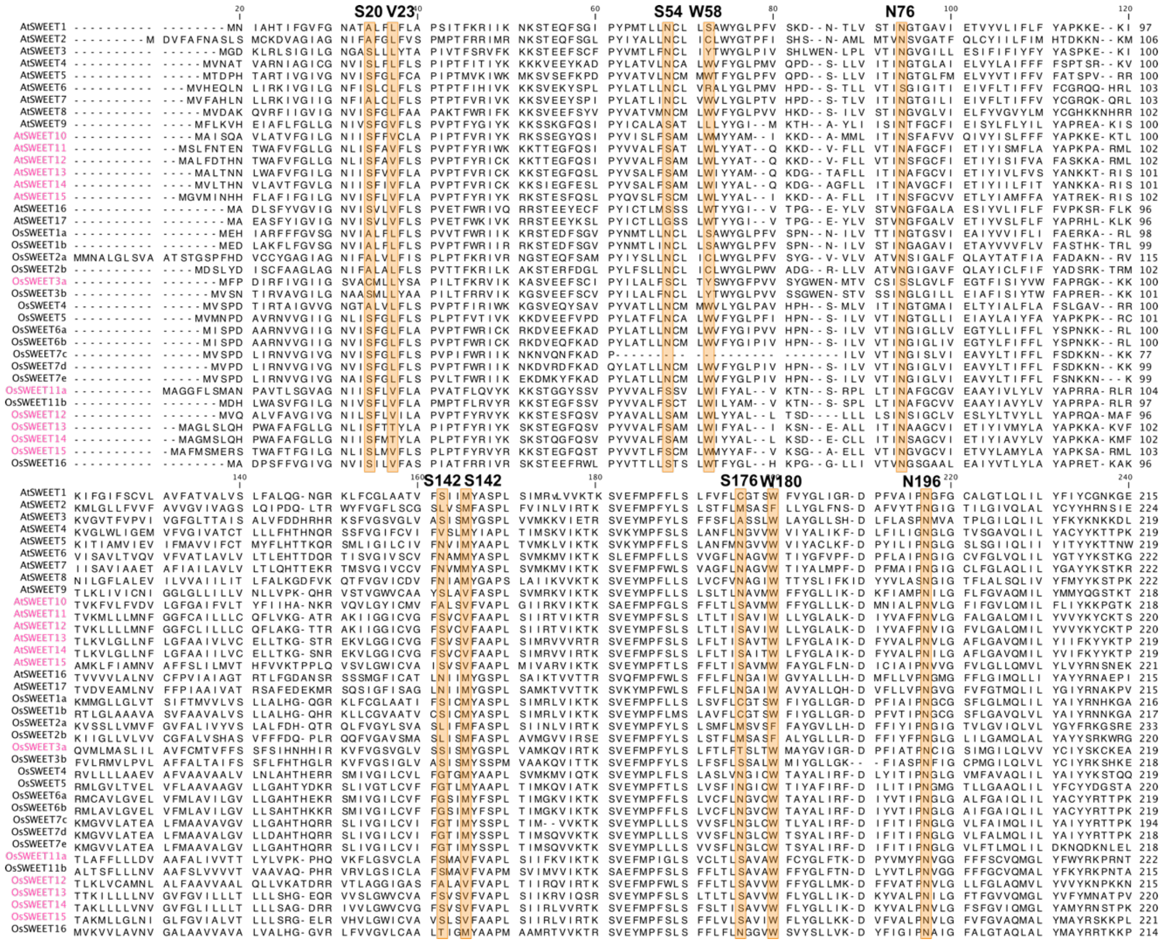

**Fig. S8. Sequence alignment of SWEET proteins from Arabidopsis and rice.**

Amino acids in the substrate binding pocket are highlighted in orange. SWEET homologs labeled in pink are able to transport $GA_3$ confirmed based on Y3H assays (1-3).

**Table S1.** Primers used in this study

| Primer | Sequence (5' > 3') |
|---|---|
| Ser20Ala-F | TTGGGTAACATCATAGCTTTCGTCGTGTTCTTGGCCCCAGTGC |
| Ser20Asn-F | TTGGGTAACATCATAAACTTCGTCGTGTTCTTGGCCCCAGTG |
| Ser20Gln-F | TTGGGTAACATCATACAATTCGTCGTGTTCTTGGCCCCAGTG |
| Ser20-R | TATGATGTTACCCAAGATTCCAAACACAAATGCCC |
| Asn76Gln-F | CTTCTCATCACCATACAAGCTTTTGGATGCGTCATCGAAACC |
| Asn76Gln-R | TATGGTGATGAGAAGAAAGGCTGTGCCATCTTTTTGC |
| Ser142Ala-F | ATTTGCGTTGGATTTGCTGTCAGTGTTTTCGCAGCTCCTTTG |
| Ser142Leu-F | ATTTGCGTTGGATTTCTTGTCAGTGTTTTCGCAGCTCCTTTGAG |
| Ser142Asn-F | ATTTGCGTTGGATTTAATGTCAGTGTTTTCGCAGCTCCTTTGAG |
| Ser142-R | AAATCCAACGCAAATCCCTCCGAGAACTTTCTCACGTGTTGAACC |
| Val145Ala-F | GGATTTTCCGTCAGTGCTTTCGCAGCTCCTTTGAGTATC |
| Val145Leu-F | GGATTTTCCGTCAGTCTTTTCGCAGCTCCTTTGAGTATCATG |
| Val145-R | ACTGACGGAAAATCCAACGCAAATCCCTCCGAGAAC |
| Asn196Ala-F | TACGTTGCCCTTCCAGCTGTATTAGGCGCGTTCTTAGGAGCTGTTCAAATG |
| Asn196Gln-F | TACGTTGCCCTTCCACAAGTATTAGGCGCGTTCTTAGGAGCTGTTCAAATG |
| Asn196-R | TGGAAGGGCAACGTAGAAGTCTTTAATAGCGAGACCGTAGAAGAGCC |
| SWEET13a-1-LP | TACGCTATGCAAAAAGATGGC |
| SWEET13a-1-RP | CGACAAAAGAAGTTGGCAAAG |
| SWEET14aLP | AAGCATCATCGATCTCAAACG |
| SWEET14aRP | CGCACCCAATATATTTGGAAG |

### Supplementary Information Text

### SI Materials and Methods

#### Missing side chain and loop modeling

The 3D model of SWEET13 from *Arabidopsis* was derived from the X-ray crystallographic structure of the SWEET13 sugar transporter of *Arabidopsis thaliana* in the inward open state, (PDB ID 5XPD) (4) (resolution 2.79 Å). The structure was generated with a SWEET13 carrying thermostabilizing mutations (V23L, S54N, V145M, and S176N) as a fusion with rubredoxin and bound to 2'-deoxycytidine 5'-monophosphate (dCMP). Rubredoxin and dCMP were removed computationally from 5XPD to create an apo SWEET13 model. The side chains of V23, S54, V145, and S176 were built using SCWRL4 (5). The missing first 5 residues of the N-terminal loop were modeled using MODELLER 9.14 (6). The model was optimized using the VTFM method applying conjugate gradients with a maximum iteration number of 300. The degree of the VTFM was set by the autosched module applying the long, thorough optimization schedule of autosched.slow. Subsequently, the model was refined by short molecular dynamics (MD) simulation (of 2 ns) and simulated annealing using the predefined function, refine.slow of refining module of MODELLER. Optimization was repeated two times with an objective function cutoff of $10^6$. Protonation states of titratable residues were determined by visual inspection and calculations with PropKa 3.1 (7).

#### Molecular docking

To construct the models of SWEET13 bound to sucrose (SWEET13*Suc) and gibberellic acid (SWEET13*$GA_3$), molecular docking of sucrose and $GA_3$ were performed using AutoDock 4.2, respectively (8). The docking sites of sucrose and $GA_3$ were assumed to be similar to that of dCMP based on the dCMP-bound SWEET13 structure (4). Docking sites were obtained by superimposition of the apo-SWEET13 model generated here onto the 5XPD structure for sucrose and GA. The centers of the grid boxes were derived by superposition of the glucosyl ring of sucrose onto the cytosine base of dCMP. The coordinates of the ether bond oxygen atom of sucrose served as the position of the grid center for both sucrose and $GA_3$. The number of grid points was chosen as 34, 34, 34 in X, Y, Z directions with a grid spacing of 0.375 Å. 150 independent docking runs were performed using the Lamarckian genetic algorithm (LGA) with a maximum number of 27,000,000 energy evaluations, the maximum number of 27,000 generations and population size of 300 for each ligand. The ligands and side chains of assumed binding residues were set to be flexible. The torsional degrees of freedom were 13 and 3 for sucrose and $GA_3$, respectively. The docking poses of each ligand were ranked by binding free energy estimations and population numbers of resulting clusters. Docking poses with the lowest binding free energy in the most populated clusters were selected as final conformations. All experimentally suggested sucrose binding residues are in contact with sucrose in its final docking pose. The protonation states of titratable residues were determined by PropKa 3.1 and visual inspection.

#### Preparation for MD simulation

A pre-equilibrated 1-palmitoyl-2-oleoyl-D-glycero-3-phosphatidylcholine (POPC) bilayer consisting of 161 lipid molecules was packed around the transporter of each system using the Membrane Builder facility of CHARMM-GUI (9, 10). The transporter was oriented with respect to the membrane in a way that charged side chains of the transmembrane domain were surrounded either by lipid head groups or water molecules. The membrane position relative to the transporter was determined based on the database of Orientations of Proteins in Membranes (OPM) (11). Each transporter-membrane system was solvated with 17,433 TIP3P (12) water molecules resulting in 80.27x80.27x129.65 $Å^3$ simulation box cells with a distance of at least 15 Å between protein surface and box face. Each box was replicated by periodic boundary conditions. Sodium and chloride ions corresponding to a physiological ion strength of 150 mM were added. Additional chloride counterions were added to achieve a neutral net charge of all systems.

The minimization procedure and set-up of MD simulations were performed with GROMACS 5.0.6 program package (13, 14) using the CHARMM-36 all-atom additive force field containing carbohydrate parameters (15). The parameters of $GA_3$ were generated by the CHARMM-GUI using the CHARMM General Force Field (CGenFF) (16). The real space summation of electrostatic interactions was truncated at 12 Å, and the Particle Mesh Ewald (PME) method was used to calculate the electrostatic interactions beyond 12 Å with a grid spacing of 1.2 Å and an interpolation order of 4. Van der Waals interactions were calculated using a cut-off of 12 Å. The solvated systems were energy minimized to eliminate unfavorable positions. Harmonic positional restraints were applied on protein, ligands, lipids, and dihedral restraints on lipid heavy atoms to achieve smooth minimization. 5,000 steps steepest descent algorithm was used adopting harmonic force constants for protein backbone/side-chain atoms -4,000/2,000 $kJmol^{-1}nm^{-2}$, for the ligand atoms - 4,000 $kJmol^{-1}nm^{-2}$, for the lipid phosphor (P) atom in Z direction (orthogonal to the membrane) - 1,000 $kJmol^{-1}nm^{-2}$, for the improper dihedral angle formed by the glycerol carbon and the oleoyl ester oxygen atoms of POPC (restricted to 120˚) and for the dihedral angle around the double bond of the oleoyl chain of POPC (restricted to 0˚) - 1,000 and 1,000 $kJmol^{-1}rad^{-2}$, respectively.

The minimized systems were equilibrated over 6 successive runs: 2 x 25 ps (*NVT*, 1 fs timestep), 25 ps (*NPT*, 1 fs timestep) and 3 x 100 ps (*NPT*, 2 fs timestep). Gradually decreasing harmonic restraints were applied to the protein, ligands, and the lipid heavy atoms. The force constant values were decreased every run according to the following procedure: the protein backbone/side-chain atoms − 4,000/2,000, 2,000/1,000, 1,000/500, 500/200, 200/50, 50/0 $kJmol^{-1}nm^{-2}$; the ligand atoms − 4,000, 2,000, 1,000, 500, 200, 50 $kJmol^{-1}nm^{-2}$; the lipid phosphor (P) atom in Z direction − 1,000, 1,000, 400, 200, 40, 0 $kJmol^{-1}nm^{-2}$; improper dihedral angle formed by the glycerol carbon and the oleoyl ester oxygen atoms of POPC (restricted to 120˚) and dihedral angle around the double bond of the oleoyl chain of POPC (restricted to 0˚) − 1,000, 400, 200, 200, 100, 0 $kJmol^{-1}rad^{-2}$.

**Production of MD trajectories**

All-atom MD simulations were performed on 96 nodes of a local computer cluster with CHARMM-36 force field using GROMACS 5.0.6 package. 2,100 ns trajectory was performed for each system (*SI Appendix,* Fig. S1). The first 100 ns of each simulation was used as non-restrained equilibration and was not included in binding pocket clustering. The following MD protocols were used: the integration time step was 2 fs; the isobaric–isothermal (*NPT*) ensemble was employed; the pressure was set to 1 bar using semi-isotropic coupling (uniform scaling of X-Y box vectors, independent Z) to the Parrinello-Rahman barostat with a time constant of 5 ps and an isothermal compressibility of $4.5*10^{-5}$ $bar^{-1}$; the temperature was kept constant at 300 K using the Nosé-Hoover thermostat with a time constant of 1 ps. Bonds with hydrogen atoms were constrained using the Linear Constraint Solver (LINCS). Atomic coordinates were recorded every 10 ps.

The MD trajectories were analyzed (root mean square deviations (RMSDs), root mean square fluctuations (RMSFs), principal component analysis (PCA), clustering, and geometric measurements) with tools included in the GROMACS 5.0.6 package. Correlation matrices were calculated with the Bio3d R package (17). The RMSDs were calculated on the backbone atoms of the transporter after the least square fitting of each snapshot on the first structure of the MD run. According to the RMSD curves, the initial 100 ns of each trajectory was omitted as non-equilibrated. To remove the overall translation and rotation of the transporter – rigid body motion – each frame of the truncated trajectory was superimposed onto the Cα atoms of the first structure of the truncated trajectory for the RMSF calculation, correlation analysis, and PCA. Only Cα atoms were considered for RMSF, correlation, and PCA computations. PCA was performed on the net conformational ensemble obtained by concatenating the individual MD trajectories of each system (6000 ns) to allow the comparison of global motions of the ligand-free and bound transporter states (18, 19). The binding pocket was defined as the ligand and the experimentally suggested (4), putative sucrose binding residues. It was assumed that $GA_3$ makes similar contacts with the transporter as sucrose, reasoning the same choice of possible binding residues. The binding site clustering was performed after superimposition of each snapshot onto the atoms of binding residues and ligands of the first equilibrated structure. The clustering was based on RMSD comparison. The Gromos algorithm of Gromacs 5.0.6 was applied for clustering analysis using a cut-off of 2 Å for both ligand-bound states. The middle structure of each cluster was considered as *representative conformation*. Noteworthy, after 900 ns - only the first most populated cluster becomes the main binding mode of sucrose (Fig. 1A).

**Sucrose transport assays in mammalian cells**

The ORF of *Arabidopsis* SWEET13 was cloned into pcDNA3.2/V5-DEST. Mutations (S20A, S20N, S20Q, V23A, S54A, N76Q, S142A, S142L, S142N, V145A, V145L, N196A, N196Q) were introduced by PCR mutagenesis using PrimeSTAR® GXL DNA Polymerase (Takara) and NEBuilder® HiFi DNA Assembly (NEB) (*SI Appendix,* Table S1). For sucrose transport assays, HEK293T cells were co-transfected with constructs carrying the sucrose sensor FLIPsuc-90µΔ1 (20) and SWEET13, or SWEET13 variants carrying mutations, by Lipofectamine LTX (Iinvitrogen) in 8-well glass-bottom chambers (Iwaki) and incubated for 48 h. Fluorescence images were acquired on a Nikon Ti2-E microscope equipped with PRIME BSI sCMOS camera (Photometrics) and SPECTRA X LED Light (Lumencor), a 40x dry objective lens CFI PlanFluor (Nikon) under excitation at 440 nm and emission channels for CFP (ET480/40m, Chroma Technology) and YFP (ET535/30m, Chroma Technology). Culture media were replaced with 150 µL Hanks Balanced Saline Salt (HBSS) buffer followed by the addition of 150 µL HBSS buffer containing 20 mM sucrose. Images were taken for 6 min at 5-sec intervals. Image quantification was performed with Fiji/ImageJ software (NIH). Data were analyzed and the box plots were generated with Prism8 (GraphPad Software).

**GA transport assays in mammalian cells**

For GA transport assays, HEK293T cells were co-transfected with constructs carrying the GA sensor GPS1 (21) and SWEET13, or SWEET13 variants carrying mutations, by Lipofectamine LTX (Invitrogen) in 8-well glass-bottom chambers (Iwaki) and incubated for 48h. Culture medium was replaced with 300 µL Dulbecco's Modified Eagle Medium (D-MEM) without phenol red containing either 0.001% (v/v) DMSO or 1.0 µM $GA_3$ dissolved in 0.001 % DMSO. Cells were incubated for 3h. Fluorescence images were acquired as for sucrose transport assays.

**GA transport assays in a Yeast three-hybrid system**

GA transport assays were performed using a previously described Y3H system (1, 3). The yeast strain PJ69-4a [MATa *trp1-901 leu2-3,112 ura3-52 his3-200 Δgal4 Δgal80* LYS2::GAL1-HIS3 GAL2-ADE2 *met2*::GAL7-lacZ] (BY5625) was obtained from the National Bio-Resource Project (NBRP, Japan). Yeast cells were cotransformed with the GA receptor components in pDEST22-GAI and pDEST32-GID1a and either pDRf1-GW (empty vector control), pDRf1-SWEET13, or pDRf1-SWEET13 with mutations by conventional lithium acetate/PEG transformation. In the pDRf1-vectors, ORF expression is driven from the strong PMA1 promoter fragment {Loque:2007ia}. Three independent colonies were used as technical replicates for each assay. Colonies were incubated in Synthetic Defined (SD -Leu, -Trp, -Ura) liquid media and incubated overnight at 30°C. Culture were diluted sequentially to 10, $10^2$, $10^3$, and $10^4$ cells/µL. 10 µL cell suspension was spotted on SD (-Leu, -Trp, -Ura) or selective (-Leu, -Trp, -Ura, -His) media containing 3 mM 3-amino-1,2,4-triazole (3-AT) in 0.001% (v/v) DMSO, and 0.1 µM $GA_3$ or 1nM $GA_4$ in 0.001% (v/v) DMSO, and incubated for 3 days at 30°C. Plates were photographed.

**Plant materials and growth conditions**

Single *sweet13* and *sweet14* mutants were obtained from the *Arabidopsis* Biological Resource Center (ABRC) and homozygous mutants were selected by PCR using primers (Table 1). The *sweet13;14* double mutant was generated by crossing *sweet13* (SALK_087791: a T-DNA is inserted before the $39^{th}$ base T of the fifth exon.) with *sweet14* (SALK_010224: a T-DNA is inserted before $16^{th}$ base A of the fifth). For complementation and other purposes, a translational fusion SWEET13-turboID-GFP (SWEET13-GFP) construct was generated from a genomic clone of SWEET13 (At5g50800), including 4059-bp upstream from translational start site and a 2178-bp region downstream from the stop codon. The turboID-GFP (22) sequence was fused to SWEET13 just before the stop codon. Using SWEET13-turboID-GFP as a template, $SWEET13^{N76Q}$ and $SWEET13^{S142N}$ mutations were introduced by inverse PCR using PrimeSTAR® GXL DNA Polymerase (Takara) and NEBuilder® HiFi DNA Assembly (NEB) (*SI Appendix,* Table S1). The turboID-GFP fused constructs were transferred to pBIN40 (23). Agrobacterium-mediated transformation by flower dipping was used to generate transgenic plants (24). Seeds were surface-sterilized with 70% (v/v) ethanol and washed with sterile water four times. After stratification at 4°C in the dark for 3 days, seeds were sown on mineral wool blocks. Plants were grown in a growth chamber at 22°C in a 16 h light / 8 h dark cycle (40 $\mu mol\ m^{-2}\ s^{-1}$ of white light) with 60% humidity for 2 weeks. Mineral wool blocks with two-week-old seedlings were transferred to soil containing vermiculite and Metro-mix at a 2:1 ratio and grown in a growth chamber at 23°C under continuous light (60 $\mu mol\ m^{-2}\ s^{-1}$ of white light). Plants were supplied with nutrients (1/1000 dilution of Hyponex) once every two weeks.

**FDA-PI staining of pollen**

To evaluate the plasma membrane integrity of pollen, pollen was stained with 1 µg/mL propidium iodide (PI) and 2 µg/ml fluorescein diacetate (FDA) in MilliQ water. After staining, pollen was observed under an LSM800 confocal microscope using a 20x dry objective Plan-APOCHROMAT (Zeiss). FDA was excited at 488 nm and detected with a window of 500-550 nm. PI was excited at 561 nm and detected with a window of 570-640 nm. Bar graphs were generated and statistical analyses were performed with Prism8 (GraphPad Software).

***In vitro* pollen germination assays**

To evaluate the pollen germination efficacy, pollen grains were incubated on pollen germination medium (0.01% boric acid, 5 mM $CaCl_2$, 5 mM KCl, 1 mM $MgSO_4$, 10% sucrose, adjusted to pH7.5 with KOH, 1.5% low-melting agarose supplemented with 10 µM epibrassinolide) at 23°C in a humid chamber (25). Images were obtained using an Axio Imager A2 upright microscope (Zeiss) equipped with a CCD camera (Axiocam 512 color; Zeiss).

**Confocal imaging of anthers**

Fluorescence images of anthers from plants stably expressing SWEET13-turboID-GFP were captured with an LSM800 confocal laser scanning microscope (Zeiss) equipped with a 20x dry objective. GFP was excited at 488 nm and detected in a window of 500-550 nm; autofluorescence was excited at 561 nm and detected in a window of 570-640 nm. For sum slices projection, 35 slices of Z-stacks in 1.5 µm intervals were taken. Fixation, clearing, and staining of anther were conducted following ClearSee protocol (26). Images were taken with a Z-stack in 1.5 µm intervals using a 40x water-immersion objective lens. GFP proteins were excited at 488 nm and detected in a window of 410-546 nm; autofluorescence was excited at 488 nm and detected in a window of 656-700 nm; CalcofluorWhite for cell wall staining was excited at 405 nm and detected in a window of 410-546 nm.

**SI References**


1. Y. Kanno, *et al.*, AtSWEET13 and AtSWEET14 regulate gibberellin-mediated physiological processes. *Nat. Commun.* **7**, 13245 (2016).

2. M. Morii, *et al.*, The dual function of OsSWEET3a as a gibberellin and glucose transporter is important for young shoot development in rice. *Plant Cell Physiol.* **61**, 1935–1945 (2020).

3. L.-B. Wu, *et al.*, OsSWEET11b, a potential sixth leaf blight susceptibility gene involved in sugar transport-dependent male fertility. *New Phytol.* **234**, 975–989 (2022).

4. L. Han, *et al.*, Molecular mechanism of substrate recognition and transport by the AtSWEET13 sugar transporter. *Proc. Natl. Acad. Sci. USA* **114**, 10089–10094 (2017).

5. G. G. Krivov, M. V. Shapovalov, R. L. Dunbrack, Improved prediction of protein side-chain conformations with SCWRL4. *Proteins* **77**, 778–795 (2009).

6. B. Webb, A. Sali, Comparative protein structure modeling using MODELLER. *Curr. Protoc. Protein Sci.* **86**, 2.9.1–2.9.37 (2016).

7. M. H. M. Olsson, C. R. Søndergaard, M. Rostkowski, J. H. Jensen, PROPKA3: Consistent treatment of internal and surface residues in empirical pKa predictions. *J. Chem. Theory Comput.* **7**, 525–537 (2011).

8. G. M. Morris, *et al.*, AutoDock4 and AutoDockTools4: Automated docking with selective receptor flexibility. *J. Comput. Chem.* **30**, 2785–2791 (2009).

9. S. Jo, T. Kim, V. G. Iyer, W. Im, CHARMM-GUI: a web-based graphical user interface for CHARMM. *J. Comput. Chem.* **29**, 1859–1865 (2008).

10. J. Lee, *et al.*, CHARMM-GUI input generator for NAMD, GROMACS, AMBER, OpenMM, and CHARMM/OpenMM simulations using the CHARMM36 additive force field. *J. Chem. Theory Comput.* **12**, 405–413 (2016).

11. M. A. Lomize, I. D. Pogozheva, H. Joo, H. I. Mosberg, A. L. Lomize, OPM database and PPM web server: resources for positioning of proteins in membranes. *Nucleic Acids Res.* **40**, D370–6 (2012).

12. W. L. Jorgensen, J. Chandrasekhar, J. D. Madura, R. W. Impey, M. L. Klein, Comparison of simple potential functions for simulating liquid water. *J. Chem. Phys.* **79**, 926 (1998).

13. H. J. C. Berendsen, D. van der Spoel, R. van Drunen, GROMACS: A message-passing parallel molecular dynamics implementation. *Comput. Phys. Commun.* **91**, 43–56 (1995).

14. M. J. Abraham, *et al.*, GROMACS: High performance molecular simulations through multi-level parallelism from laptops to supercomputers. *SoftwareX* **1-2**, 19–25 (2015).

15. J. Huang, A. D. MacKerell, CHARMM36 all-atom additive protein force field: validation based on comparison to NMR data. *J. Comput. Chem.* **34**, 2135–2145 (2013).

16. K. Vanommeslaeghe, *et al.*, CHARMM general force field: A force field for drug-like molecules compatible with the CHARMM all-atom additive biological force fields. *J. Comput. Chem.* **31**, 671–690 (2010).

17. B. J. Grant, A. P. C. Rodrigues, K. M. ElSawy, J. A. McCammon, L. S. D. Caves, Bio3d: an R package for the comparative analysis of protein structures. *Bioinformatics* **22**, 2695–2696 (2006).

18. M. J. M. Niesen, S. Bhattacharya, N. Vaidehi, The role of conformational ensembles in ligand recognition in G-protein coupled receptors. *J. Am. Chem. Soc.* **133**, 13197–13204 (2011).

19. M. Grosso, A. Kalstein, G. Parisi, A. E. Roitberg, S. Fernandez-Alberti, On the analysis and comparison of conformer-specific essential dynamics upon ligand binding to a protein. *J. Chem. Phys.* **142**, 245101 (2015).

20. I. Lager, L. L. Looger, M. Hilpert, S. Lalonde, W. B. Frommer, Conversion of a putative Agrobacterium sugar-binding protein into a FRET sensor with high selectivity for sucrose. *J. Bio. Chem.* **281**, 30875–30883 (2006).

21. A. Rizza, A. Walia, V. Lanquar, W. B. Frommer, A. M. Jones, In vivo gibberellin gradients visualized in rapidly elongating tissues. *Nat. Plants* **3**, 803–813 (2017).

22. T. C. Branon, *et al.*, Efficient proximity labeling in living cells and organisms with TurboID. *Nat. Biotechnol.* **36**, 880–887 (2018).

23. R. J. Iwatate, *et al.*, Covalent self-labeling of tagged proteins with chemical fluorescent dyes in BY-2 cells and Arabidopsis Seedlings. *Plant Cell* **32**, 3081–3094 (2020).

24. S. J. Clough, A. F. Bent, Floral dip: a simplified method for Agrobacterium-mediated transformation of *Arabidopsis thaliana*. *Plant J.* **16**, 735–743 (1998).

25. K. Muro, *et al.*, ANTH domain-containing proteins are required for the pollen tube plasma membrane integrity via recycling ANXUR kinases. *Commun. Biol.* **1**, 152 (2018).

26. D. Kurihara, Y. Mizuta, Y. Sato, T. Higashiyama, ClearSee: a rapid optical clearing reagent for whole-plant fluorescence imaging. *Development* **142**, 4168–4179 (2015).
